# Evidence for different greater-persistence strategies under lower and higher challenge for alcohol in female rats

**DOI:** 10.1101/2022.05.18.492488

**Authors:** Thatiane De Oliveira Sergio, David Darevsky, Vanessa de Paula Soares, Maryelle de Cassia Albino, Danielle Maulucci, Sarah Wean, Frederic W. Hopf

## Abstract

Problem alcohol drinking is a substantial social and economic burden. Studies show that the misuse of alcohol is increasing in women, and that women can face higher consequences from alcohol exposure, but females have historically been understudied. Thus, there is considerable interest in understanding potential sex-different and -similar cognitive/emotional strategies, and underlying mechanisms, for alcohol responding, which would inform more effective, personalized treatments. Here, we used large cohorts of adult Wistar rats (28 females, 30 males) to provide robust assessment of potential sex differences in responding for alcohol-only and under compulsion-like drinking with moderate or higher challenge (since intake despite negative consequences can be a major obstacle to human treatment). Females had similar total licking but higher intake for all drinking conditions. However, females had significantly longer bouts under alcohol-only and moderate challenge, but not higher challenge. Further, under higher challenge, females retained several aspects of responding not seen in males, including more efficient lick volume and earlier onset of longer bouts. In addition, females overall licked slightly faster, but licking speed averaged within-bout showed no sex differences, and female intake level under alcohol-only and moderate challenge was unlinked from licking speed (unlike males, where slower licking predicted lower intake). We interpret these differences as greater persistence-like responding but not vigor per se in females, and with different strategies under lower versus higher challenge. Finally, drinking levels did not differ across the estrous cycle, although ovariectomy reduced alcohol-only and moderate-challenge intake. Together, while many aspects were sex-similar, suggesting some common drinking mechanisms, there was clear evidence for (perhaps more nuanced) sex-different alcohol strategies, which might make an outsized contribution to excessive drinking since women can have more drinking problems. Thus, our studies provide important context for future work examining sex differences in pathological drinking mechanisms.

## INTRODUCTION

Alcohol Use Disorder (AUD) is among the most prevalent mental disorders [1, 2], extracting ∼$250 billion/year in the US alone and producing a number of medical and social harms [3–10]. One particular concern is that problem alcohol drinking in women has risen dramatically in recent years, while rates in men have remained stable or declined [1,11–15]. Importantly, in addition to rising intake, women can have greater alcohol problems including negative impacts on health [16–19] and may develop dependence faster [20–22]. However, female subjects have historically been excluded from many studies, and only in recent times have sex differences in underlying mechanisms been more widely investigated. Such differences are important to consider when developing personalized treatments for addiction and related conditions [23–25], since reducing intake can decrease health risks and relapse [8–10], but at present there are limited treatment options for AUD [26].

In addition to alcohol-only drinking (AOnly), we and others have focused in particular on compulsion-like alcohol drinking (CLAD), where intake persists despite negative consequences, since this can be a major obstacle to treatment and a strong driver of intake and its substantial harm [27–36]. Intake despite negative consequences features prominently in the DSM-V definition of AUD (e.g. [37, 38]), and several groups find less sensitivity to cost with AUD [39–41] (but [42, 43], see Discussion), while greater drinking is associated with more alcohol problems [38,44–46]. Thus, it is critical to understand potentials sex differences in the behavioral expression and mechanisms of consequence-resistant drinking as well as alcohol-only intake, especially since women with AUD can have greater alcohol problems.

While human clinical challenges propel the search for clearer indicators to help guide improvement of treatment, rodent models can provide valuable direction by allowing mechanistic investigations. More generally, willingness of rodents to respond for reward paired with aversive consequences is considered to model some aspects of compulsion-like responding in humans [31,32,35,36,47]. In addition, human [48, 49] and rodent [50–53] findings converge upon the common importance of particular cortico-striatal connections (including insula) for compulsion-like responding for alcohol, and these circuits are likely critical for many other addiction-related behaviors [54]. Sex differences are known to exist for several addictive behaviors in rodents, e.g. where female often drink more alcohol than males [18, 25]. However, much remains to be learned about sex-different and sex-general behavioral patterns of responding for alcohol, which can provide indications of potential sex differences in strategy when responding for alcohol, and is a critical foundational step to help discover mechanisms underlying sex differences. Another concern is what has been come to be known as the replicability crisis, where different researchers can find divergent results.

To address these different challenges, we set out to compare sex-related patterns of responding for alcohol in two large cohorts, examined in the states of California and Indiana, with a total of n=28 female and n=30 male adult Wistar rats. Each rat drank under conditions of alcohol-only, moderate-challenge (10 mg/L quinine in alcohol, AQ10) and higher-challenge compulsion-like intake conditions (60 or 100 mg/L quinine in alcohol, AQ60 or AQ100), generating a very large data pool to investigate potential sex differences in responding for alcohol. Also, our studies utilize lickometry to study the behavioral microstructure of licking, which is widely used to infer motivation (see [55, 56], and can allow us to gain deeper insight into potential underlying action strategies and sex differences therein. We previously utilized this paradigm in male rats to discover that CLAD under moderate challenge involves decreased response variability [47, 55], agreeing with the suggested importance of automaticity for compulsion [35,36,57,58]. These findings also perhaps reflect a behavioral strategy which allows action while minimizing need to attend to and be impacted by adverse consequences (which we call the “head down and push” model of aversion-resistant responding).

In the present study, we demonstrate that females showed several behavioral patterns indicating greater persistence in responding for alcohol relative to males. Females had significantly longer bouts under alcohol-only and moderate-challenge CLAD, which was lost under higher challenge. However, under higher challenge, females retained several behaviors which were impaired in males, including greater intake relative to the number of licks, and the ability to generate longer bouts earlier in the session. Faster licking has been linked to greater motivation and vigor (see [47, 55]), but the noted differences in female responding were not accompanied by changes in average licking speed, and female intake level under alcohol-only and moderate challenge were unlinked from licking speed, unlike males where slower licking predicted lower intake. Finally, we find no impact of estrous cycle on female drinking. Together, we interpret these sex-related differences as greater persistence-like responding in females without changes in vigor per se, and with two different persistence-like patterns in females versus males under lower versus higher challenge.

## METHODS

### Alcohol drinking

All experiments conducted following NIH guidelines and approved by UCSF and Indiana University IACUCs. Rats arrived at PN45-50 age and were single housed in clear plastic cages in housing rooms with approximately equal numbers of females and males, with dark cycle 1130am-1130pm. After ∼2wk habituation, alcohol drinking began. Our data come from two large female/male cohorts, one performed in California (CA, n=15 females and 15 males) and one in Indiana (IN, n=13 females and 15 males), in part to address overall issues raised in relation to the “replicability crisis.”

Rats initially drank under intermittent access two-bottle-choice (20% alcohol in water, vs water-only) (IA2BC), with 16-24-hr intake each day starting Monday, Wednesday and Friday at ∼1hr into the dark cycle. After >3 mo IA2BC, rats were switched to limited daily access (LDA), with intake 20min/d, 5d/wk, primarily to increase number of experiments per rat. After 3-4wk LDA, rats had 2-3 days/wk where they drank quinine in alcohol, typically 2 sessions AQ10, then 2 AQ60, then 2 AQ100. We typically do this quinine habituation [47,50,51,55,59–61], in part to assure that AQ is not novel during actual experimental sessions.

Experimental sessions used lickometers, and, since rats primarily drink at the onset of the LDA session (as we address in [47, 55]), sessions were 5 min long (as we used in [59]). Lickometers were custom made, with custom-written C++ code running on an Arduino Uno with an Adafruit MPR121 capacitive touch sensing breakout board to record lickometry data; a very small capacitive current is run through the 7.6 cm metal licking tube, and the discharge is detected when the rat’s tongue contacts the metal tube. We have used lickometry to search for potential behavioral changes indicating use of different action strategies during different aspects of alcohol drinking [47, 55], and this capacitative type of lickometer system has previously been used e.g. for in vivo electrophysiology studies addressing brain mechanisms of licking [62–65].

Importantly, from each rat, we collected two sessions from each of four different drinking conditions: AOnly, alcohol with moderate challenge (AQ10), alcohol with higher challenge (AQ60 or AQ100) [47, 55]. The four drinking conditions were each run once, in randomized order within a given rat, and then run a second time, as we have done [47,50,51,55,59–61]. Thus, with 28 female and 30 male rats, this gives 56 sessions for each drinking condition for females and 60 sessions per condition for males. We note that one Fem Q10 lickometer file was lost, although we keep this data point when comparing g/kg, total lick, and lick volume measures. In addition, as shown in **Suppl. Fig.1**, a very small number of sessions showed quite high g/kg relative to total licks, which could suggest the tongue stayed in contact with the sipper tube (and not registering separate licks). There were also sessions with very few licks (<10) which had no bout. These were all removed from analyses of session-level lick timing and patterns, with the number of such excluded sessions 5, 4, 6, 1 for AOnly, AQ10, AQ60, and AQ100 for females, and 2, 3, 1, 3 for males. Thus, session-level analyses still had >50 for each drinking condition, but the main findings persist (with a few shifting to trend level with multiple corrections) when including all sessions (see **Suppl. Fig.1**)

We also note that rats examined in IN rats drank significantly more than CA rats (detailed in **Suppl. Table1**). However, sex differences were largely conserved across the CA and IN cohorts, e.g. with females having longer AOnly and AQ10 bouts than males (MW for sex differences; ave bout length by bout: AOnly CA *p=*0.0359, AOnly IN *p=*0.0426, AQ10 CA *p=*0.0065, AQ10 IN *p=*0.0480; ave bout length by session: AOnly CA *p=*0.0373, AOnly IN *p=*0.0097, AQ10 CA *p=*0.0035, AQ10 IN *p=*0.0558).

For the study of the estrous cycle, we used 30 females rats from IN and 15 females from CA, some of which overlapped with rats used in sex difference analyses. All the OVX rats used on the study were tested only in IN.

### Analyses of responding for alcohol

Lickometry analyses were performed as we previously published [47, 55]. We examined licking which was aggregated into bouts, where a bout is defined as 3 licks with <1sec between licks, which we previously used based on extensive previous literature (discussed in [55]). Licking speed was assessed by determining the time between each pair of licks, the Inter Lick Interval (ILI). Bout and other lickometry measures were determined using custom-written programs in Python.

Also, as we have done before [47, 55], and following other groups [66, 67], we analyzed responding across sessions or bouts separate from the animal identities; Darevsky and Hopf (2020) has a detailed validation of such analyses. For analyses by bout, we determined (for each bout) the bout length, start time of the bout, average licking speed, standard deviation of licking speed, and initial licking speed (average of the first two ILIs of the bout). Analyses by session had a greater range of measures, including g/kg intake, total licks, longest bout, averaged bout length, number of bouts, averaged bout licking speed and standard deviation, stray licks (licks outside of bouts), lick volume (average alcohol volume per lick), and g/kg alcohol divided by total licks (another measure of average intake level per lick).

### Assessment of quinine sensitivity in alcohol or water

To assess how quinine in alcohol could reduce intake level, we normalized the alcohol-quinine (AQ) drinking level in a given session relative to the related AOnly session and expressed this relation as log[100*(AQ-intake/AOnly-intake)]. We have used this method before [59–61], and briefly, as detailed in Lei et al. (2019), it reduces the impact of outliers, relative to analyses determining percent change in consumption, e.g. where percent change is more influenced by lower AOnly intake values. Since the Y axis is a log scale, Fig.1C has dotted red lines to indicate where CLAD intake was 0%, 50%, or 80% lower than AOD.

**Figure 1.**
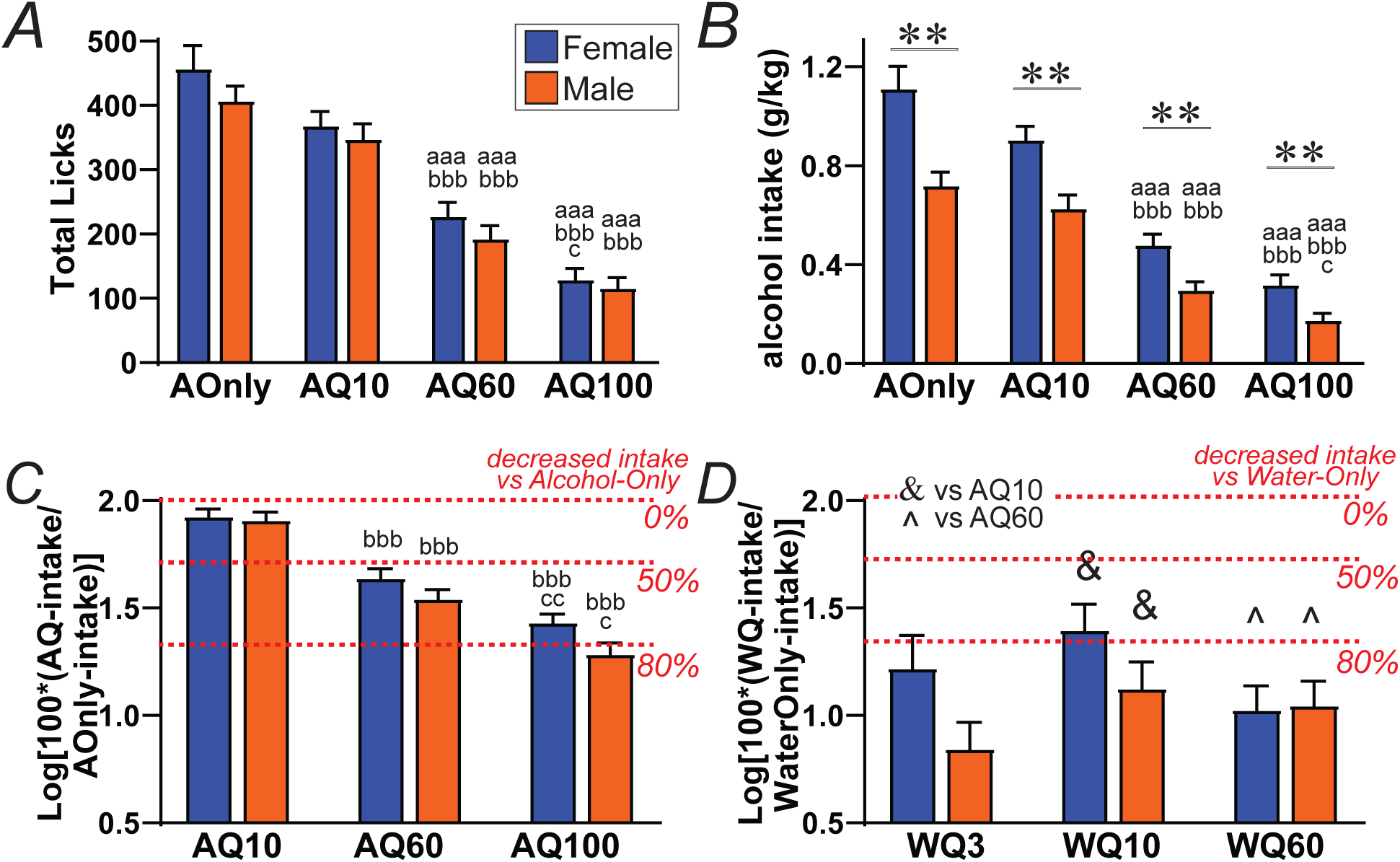
Equivalent aversion-resistance in female and male rats. **(A)** Sex-similar total licking at each drinking condition, and similar reduced licking under higher challenge. **(B)** Greater alcohol intake level in females at all drinking conditions. **(C)** Sex-similar decreases in alcohol intake with higher quinine levels. **(D)** Quinine 3, 10, or 60 mg/L in water similarly and significantly (>80%) reduced water intake in females and males. **(C, D)** are in a log scale, and dotted lines show 50% and 80% decrease in intake (see Methods). ∗∗ *p<*0.0025 MW effect of sex. For letters: aaa *p<*0.001 effect of drinking condition vs AOD (KruskalWallis with Dunn’s post-hoc); bbb *p<*0.001 AQ60 or AQ100 vs AQ10; c *p<*0.05 Q100 vs Q60. & *p<*0.001 WQ10 vs AQ10, ^ *p<*0.001 WQ60 vs AQ60.

To assure that AQ intake reflected aversion-resistance, and to assess basic sensitivity to quinine aversion in females and males, we determined intake of Water-Only (WOnly) versus water adulterated with 3, 10, or 60 mg/L quinine (WQ3, WQ10, WQ60, n=14 of each sex, a different cohort from alcohol-drinking experiments). W±Q intake was measured in 20min/d 5d/wk sessions ∼3hr before alcohol drinking each day. Rats had ∼3wk WOnly training before WQ sessions began. WOnly intake was more variable than alcohol, thus baseline WOnly for each rat was the average of six sessions. We then performed 3 sessions each of WQ3, WQ10, and WQ60 per rat, which were examined independent of rat identity (see above) and normalized to basal WOnly with log[100*(WQ-intake/WOnly-intake)].

### Estrous cycle analysis

To assess whether the phases of the estrous cycle influence alcohol drinking, we collect the vaginal epithelium of intact female rats. Before starting the collection of the vaginal smear, the rats were handled for 5 days prior to get used to the experimenter. During the collection, the female was gently hold and the tip of a sterile pipette with 10µl of sterile saline (NaCl 0.9%) was inserted into the rat’s vagina, but not deeply, in order to avoid cervical stimulation and pseudopregnancy. The liquid was flushed into the vagina and the fluid placed on a glass slide for post-analysis. The collection happened once a day after the drinking sessions. The slides were stained (Differential Quick Staining Kit – Electron Microscopy Services), and pictures of the estrous cycle were analyzed by two blind experimenters. The phases of the estrous cycle were defined based on Cora and colleagues (2015 [68]), with sample images and criterion for different stages shown in **Suppl. Fig.7**. We note that we have fewer proestrus samples, which may reflect where proestrus averages 14hrs in rats and usually occurs in midmorning, and our tests were carried during the dark phase (usually 1-2hrs after light goes off), so our females were perhaps more likely to be transiting to estrous [68].

### Ovariectomy

Female rats had both ovaries removed. After the surgery, animals were monitored every day, and after one week returned to alcohol intake. Behavioral tests were conducted between 5 – 6 months after the surgery.

### Statistical Comparisons

Most of our response data was not normal or log normal (determined with Shapiro-Wilk test). Thus, we predominantly used non-parametric statistics. First, to compare across drinking conditions within a given sex, we used Kruskal-Wallis (KW) followed by Dunn’s post-hoc to examine pair-wise across the drinking conditions within that sex. Second, to compare across sexes for a given drinking condition (e.g. total licks during alcohol-only intake in females versus males), we used Mann Whitney (MW). Statistics were performed with GraphPad Prism. It is also important to note that we report uncorrected p values. However, the p value required to indicate significance for a given intake measure was adjusted for multiple comparisons. For MW assessment for a given intake measure, we divided *p<*0.05 by 4 (responding compared across sexes for the 4 drinking conditions), giving *p<*0.0125 to be considered significant. For KW, Dunn’s posthoc p values are already corrected for multiple comparisons by Prism.

## RESULTS

One primary goal was to have a robust comparison of female and male alcohol drinking patterns, to identify behavioral indicators of potential sex differences in response strategies. Briefly, as detailed in Methods, data are combined from two large female/male cohorts, and, within each rat, we examined intake and licking patterns in two sessions from each of the four different drinking conditions: AOnly, alcohol with moderate challenge (i.e., AQ10), and alcohol with higher challenge (AQ60 or AQ100), tested in randomized order. Also, as described in Methods, and as we [47, 55] and others [66, 67] have done, we analyzed responding across sessions or bouts separate from the animal identities.

### Females have similar number of licks and greater intake levels

Females and males were very similar in the total number of licks exerted for alcohol, with no sex differences in licking for any drinking condition (**Fig.1A**; MW, all *ps>*0.19). Further, as quinine level increased, females and males both decreased total licking (KW, *p<*0.0001 in each sex). Post-hoc analyses showed that, while there was no difference between AOnly and AQ10 licking in either sex (*ps>*0.7), responding was significantly reduced under AQ60 and AQ100 relative to AOnly and AQ10 in both females and males (all *ps<*0.001). Thus, females and males had a similar number of licks for each drinking condition, and equivalent decreases in licking under higher challenge.

In contrast to total licks, the total alcohol intake (g/kg) was significantly greater in females versus males for all drinking conditions (**Fig.1B**; MW, all *ps<*0.001), likely related to smaller body size (females: 363±10 g; males 621±15 g). In addition, similar to total licks, AQ10 drinking levels were not different from AOnly in either sex (*ps>*0.9), but higher quinine (60 and 100 mg/L) significantly decreased alcohol intake level compared to AOnly or AQ10 in both sexes (all *ps<*0.001). These findings agree with our previous studies [47, 55] that male rats persist in drinking AQ10 (and thus are considered “aversion-resistant under moderate challenge”), while Q60 and Q100 reduced intake when added to alcohol.

To better assess whether there were sex differences in quinine effects on alcohol consumption, we first expressed the intake level under different CLAD conditions relative to drinking level under AOnly (**Fig.1C**), using log[100*(AQ-intake/AOnly-intake)] (see Methods). Since Y-axis is a log scale, dotted red lines in **Fig.1C** indicate where CLAD intake was 0%, 50%, or 80% lower than AOD. In agreement with **Fig.1B**, normalized intake under AQ60 and AQ100 was significantly lower than AQ10 in both sexes (KW, *ps<*0.0001). There were no significant sex differences in AQ10÷AOnly (MW *p=*0.9791), AQ60÷AOnly (*p=*0.3086), and AQ100÷AOnly (*p=*0.0195, n.s. with multiple corrections). Thus, female and male alcohol intake was similarly impacted in a dose-dependent manner by quinine (with a trend for females to drink more AQ100), even while female absolute intake levels were higher than males under all drinking conditions (**Fig.1B**).

### Females and males have similar sensitivity to quinine in alcohol

One critical question is whether females and males differed in their basic aversion sensitivity. Thus, in a separate cohort of rats, we examined how water intake would be impacted by addition of 3, 10, or 60 mg/L quinine (**Fig.1D**). Unlike AQ, there were no significant differences in water intake level with 3, 10, or 60 mg/L quin (KW female *p=*0.1008, male *p=*0.2014), indicating that water intake was reduced by >80% by all quinine doses tested. There were also no sex differences in sensitivity to quinine in water (MW: WQ3 *p=*0.0885; WQ10 *p=*0.1350; WQ60 *p=*0.8334). With these data, we were also able to compare the impact of a given dose of quinine on alcohol vs water intake. Importantly, the impact of 10 mg/L quin on water was significantly greater than the effect of 10 mg/L quin in alcohol in both sexes (MW, *ps<*0.001), with similar findings with 60 mg/L quin in water versus alcohol (MW, *ps<*0.001). These findings provide important confirmation that females and males were both aversion-resistant under AQ10 and AQ60 conditions, and also that there were no sex differences in basic sensitivity to quinine.

### Females have significantly longer and fewer bouts except under higher challenge

While females had very similar total licks as males for each drinking condition, more detailed analyses of lick microstructure indicated sex differences in response patterns. Importantly, females had significantly longer bouts than males under AOnly and AQ10, which was lost under AQ60 and AQ100. Thus, the longest bout within a given session was significantly greater in females than males for AOnly (MW=0.0001) and AQ10 (MW=0.0031) but showed no sex differences for AQ60 (*p=*0.2029) or AQ100 (*p=*0.2678) (**Fig.2A**). Similarly, the average bout length within a session was significantly longer in females for AOnly (MW *p=*0.0018) and AQ10 (*p=*0.0013), with no sex differences for AQ60 (*p=*0.3560) or AQ100 (*p=*0.0511) (**Fig.2B**). When bout lengths were examined as individual bouts, female bouts were significantly longer than males for AOnly (MW *p=*0.0013) and AQ10 (*p=*0.0010), with no sex differences for AQ60 (*p=*0.1353) although a trend for AQ100 (*p=*0.0226 n.s. with multiple comparisons) (**Fig.2C**). For all such measures, bout length shortened significantly for higher challenge (KW *p<*0.0001 for both sexes), with AOnly significantly different from AQ60 and AQ100 (*ps<*0.001) but not AQ10 (*ps>*0.9). Further, if female bouts were longer under AOnly and AQ10, with no differences in total licks, one would expect fewer bouts (**Fig.2D**). Indeed, females had significantly fewer bouts than males under AOnly (MW *p=*0.0013) and AQ10 (MW *p=*0.0009), but not AQ60 (MW *p=*0.5323) or AQ100 (MW *p=*0.8490). Also, there were no differences in bout length across drinking conditions in females (KW *p=*0.2504) but there were in males (KW *p<*0.0001). Thus, females had 40-50% longer bouts than males under AOnly and AQ10, which was lost under the higher challenge conditions, suggesting that females utilized a different response strategy than males under AOnly and AQ10; we interpret this, along with other behavioral changes (below), as an increase in persistence-like responding for alcohol in females.

**Figure 2.**
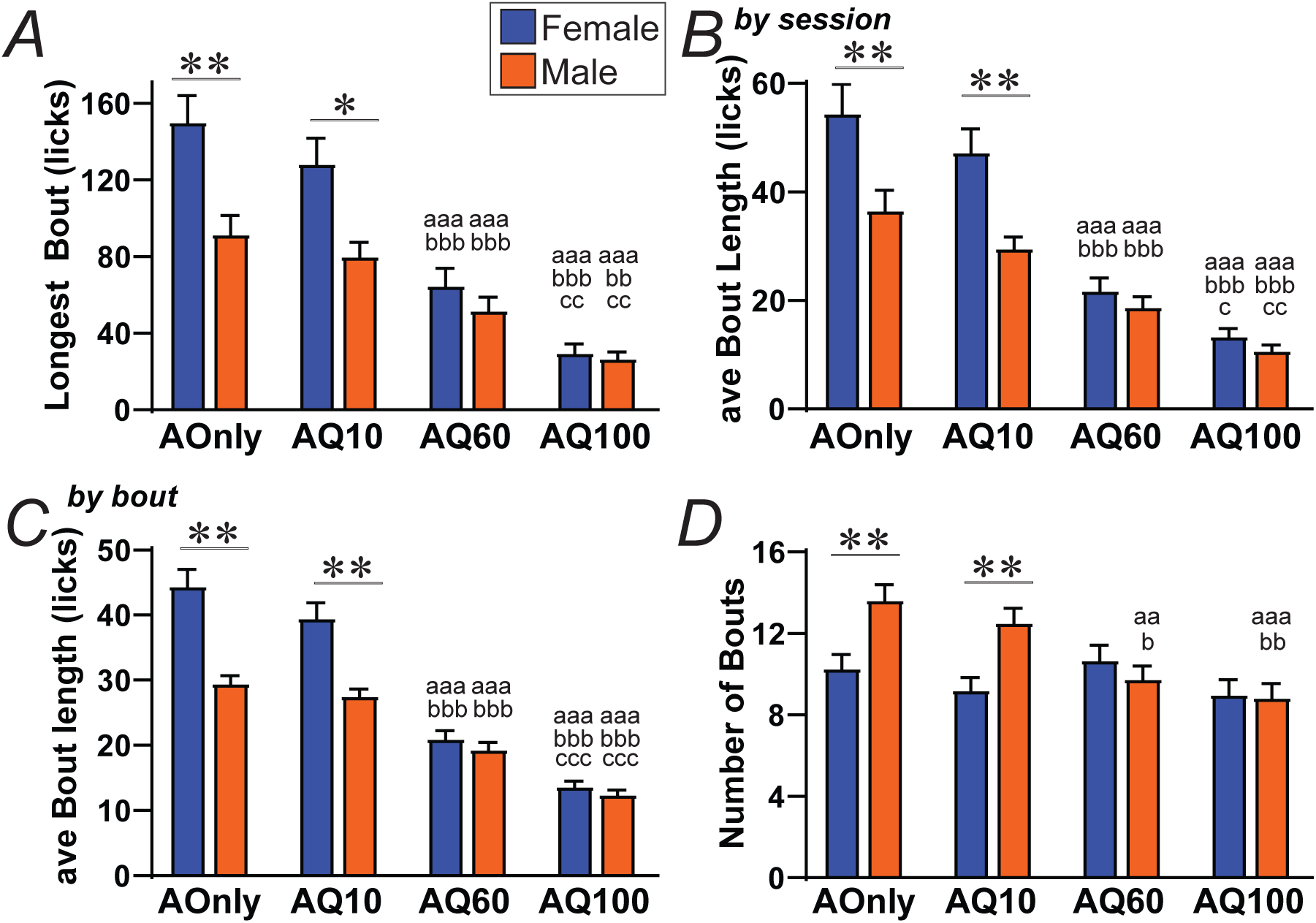
Females have longer and fewer bouts under AOnly and AQ10. **(A)** Longest bout in a session and **(B)** average bout length within a session were significantly longer in females versus males for AOnly and AQ10, but not AQ60 and AQ100. **(C)** Average bout length (averaged by bout) was significantly longer in females only for AOnly and AQ10. **(D)** Females had fewer bouts for AOnly and AQ10 but not AQ60 or AQ100. ∗,∗∗ *p<*0.0125, *p<*0.0025, MW effect of sex. For letters: aaa *p<*0.001 effect of drinking condition vs AOD; bb, bbb *p<*0.01, *p<*0.001 AQ60 or AQ100 vs AQ10; c, cc, ccc *p<*0.05, *p<*0.01, *p<*0.001 Q100 vs Q60.

### Females have significantly greater g/kg per lick under higher but not lower challenge

Since females had significantly longer bouts under AOnly and AQ10 but not higher challenge, we next looked for sex differences only under higher challenge conditions. Interestingly, we found such a difference when comparing the relation between g/kg and total licking in a session. Overall, g/kg was highly correlated with total licks in all sex/drinking conditions (all *ps<*0.0001, with R^2^ > 0.58) (e.g. similar to our studies, [47, 55]). Interestingly, the slope for this g/kg-vs-total lick relation was not sex-different for AOnly (**Fig.3A** *p=*0.1386) or AQ10 (**Fig.3B** *p=*0.2388), but were significantly different between females and males for AQ60 (**Fig.3C** *p<*0.0001) and AQ100 (**Fig.3D** *p<*0.0001). Further, under higher challenge, the slopes (licks per g/kg) were much steeper in males but were not different in females across drinking conditions (**Fig.3E**). This suggested that, under higher challenge, females were able to maintain the licking-intake relationship, while males were not.

**Figure 3.**
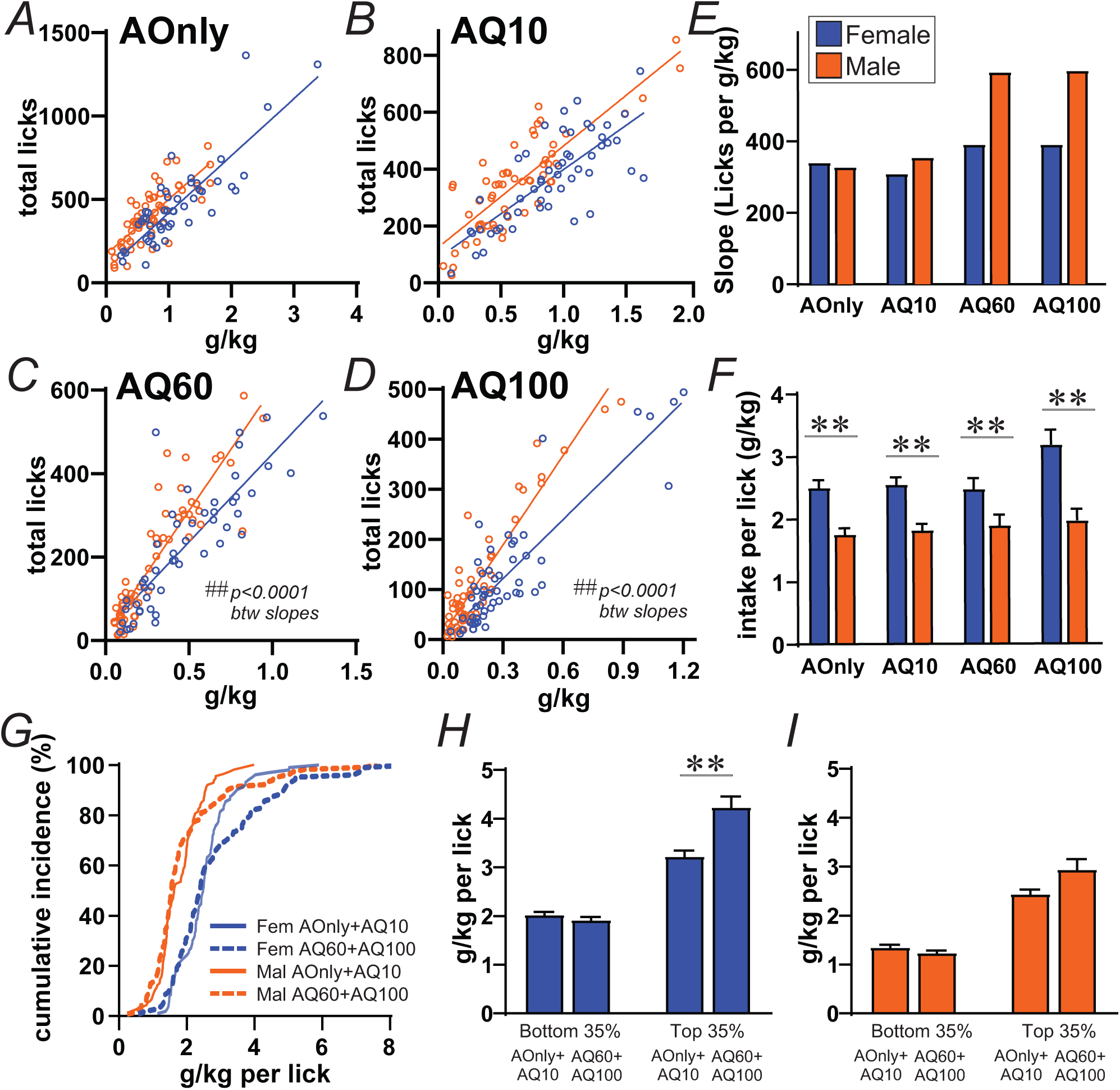
Females have larger volume per lick in some AQ60/AQ100 sessions. **(A-D)** Relation between alcohol intake (g/kg) and total licks, with no sex differences for AOnly **(A)** and AQ10 **(B)**, but significant sex differences for AQ60 **(C)** and AQ100 **(D). (E)** The slope of Licks/(g/kg) was higher for males for AQ60 and AQ100, but did not differ across drinking conditions for females. **(F)** On average, females have greater g/kg intake per lick for all drinking conditions. **(G)** Cumulative incidence of g/kg-per-lick across AOnly+AQ10 versus AQ60+AQ100 sessions. **(H,I)** The cumulative distributions in **(G)** show divergences in females at ∼65% of sessions, and the g/kg-per-lick for the top 35% of sessions was significantly greater in higher challenge vs AOnly+AOnly in females **(H)** but not males **(I)**. ∗∗ *p<*0.0025, MW effect of sex; ## *p<*0.0001 for correlation.

Thus, we next g/kg divided by total licks [g/kg-per-lick], as an indicator of lick volume. Overall, females had significantly greater g/kg-per-lick across all drinking conditions (all *ps<*0.0001) (**Fig.3F**), suggesting strong overall sex differences in lick volume efficiency, largely separate from drinking condition. There were no differences across drinking conditions in either sex for g/kg-per-lick (KW female *p=*0.0791, male *p=*0.5864), suggesting overall that this assessment of intake volume per lick was a more basic feature of licking. Measures of lick volume per se (ul alcohol/lick) were somewhat more complex, with females having significantly lower lick volume for AOnly (*p=*0.0008) and AQ60 (*p=*0.0004), males having greater for AQ10 (*p=*0.0010), and no differences for AQ100 (*p=*0.9296), and lick volume was not different across drinking conditions in either sex (KW female *p=*0.0198 with no signif post-hocs, male *p=*0.1986) (**Supp.Fig.2**); this may be complicated by differing body sizes (see below), and thus we focused on g/kg per lick, since this accounts for smaller body size in females.

When examining the slope relationships in **Fig.3A-D**, we noted that there might be shifts in lick volume in only a subset of sessions, which change the overall shape of the distribution. To simplify assessment of this possibility, we combined data from AOnly and AQ10 into a “lower challenge” group, and AQ60 and AQ100 into a “higher challenge” group. The cumulative distributions shown in **Fig.3G** demonstrate that a subset of sessions under higher challenge did show higher g/kg-per-lick. Since the divergence point was at ∼65% of sessions, we the separated g/kg-per-lick values below and above this point. The g/kg-per-lick values in the lower 65% of sessions was not different between lower and higher challenge condition in females (MW *p=*0.1831) or males (*p=*0.0870) (**Fig.3H**). However, in females, the g/kg-per-lick values in upper 35% of sessions was significantly greater in higher versus lower challenge groups (*p<*0.0001), while there was no difference in males (*p=*0.5329) (**Fig.3I**). These findings support the possibility that females have a shift in response strategy in some higher-challenge sessions, which helped to retain higher intake levels while executing fewer licks.

### Females have limited changes in measures of bout timing

Changes in licking speed measures are often observed and taken to indicate altered drive for the reward, e.g. where rats lick faster for higher sucrose (e.g. [56]), where quicker responding can indicate vigor for action [69], and where we found that male rats lick more slowly under higher challenge for alcohol, in concert with reduced intake [47, 55]. Thus, we examined inter-lick interval (ILI, the time between successive licks) measures to assess licking speed. Also note that, like many measures examined, the distribution of data was highly non-normal, for cases where differences are not be apparent in averages, and thus we also shown median, where differences are more apparent.

We first examined licking speed across all ILIs, separate from bout or session, and looked separately at ILIs within bouts, and ILIs outside bouts (which we call Inter-Bout Intervals). Within bouts, females showed a small but significantly faster responding under all drinking conditions (**Fig.4A,B**, MW all *ps<*0.0001). There were also significant differences across drinking condition in both sexes (KW *ps<*0.0001), with slower licking under higher challenge (see Fig. legend); histograms of ILI distributions are shown in **Suppl. Fig.3**. Females had significantly faster Inter-Bout Intervals under AOnly (MW *p<*0.0001), AQ10 (*p=*0.0011), and AQ60 (*p=*0.0003), with a trend for AQ100 (*p=*0.0348) (**Fig.4C,D**). Unlike within-bout ILIs, when examined across drinking conditions (KW female *p<*0.0001, male *p=*0.0012), AOnly and AQ10 were different from AQ100 (*ps<*0.007) but not from AQ60 (*ps>*0.39) in both sexes. Thus, females had slightly but significantly faster responses overall, including slowed responding within higher challenge bouts (as we previously showed in males [47]). However, Inter-Bout timing (perhaps reflecting willingness to re-engage in a bout) was only significant disrupted in both sexes under AQ100.

**Figure 4.**
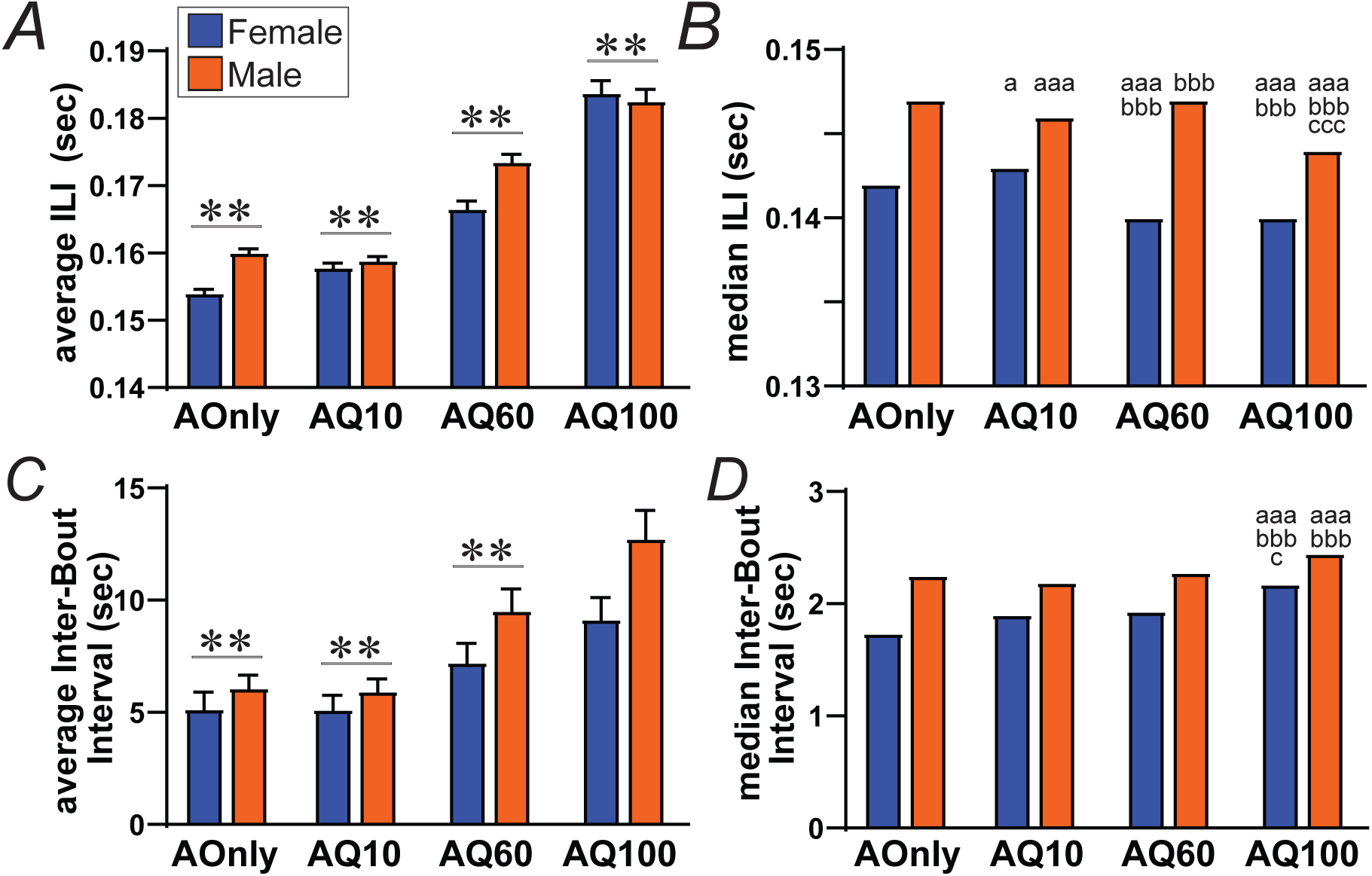
Females have faster overall licking. **(A,B)** Average **(A)** and median **(B)** of ILIs, taken as individual ILIs that occurred within bouts. **(C,D)** Average **(C)** and median **(D)** of Inter-Bout Intervals (ILIs that occurred outside bouts). ∗∗ *p<*0.0025, MW effect of sex. For letters: a, aaa *p<*0.05, *p<*0.001 effect of drinking condition vs AOD; bbb *p<*0.001 AQ60 or AQ100 vs AQ10; c, ccc *p<*0.05, *p<*0.001 Q100 vs Q60.

We next examined licking speed where we averaged the ILIs within each bout, and, for the session measure, averaged bout licking speed across all bouts within the session. When analyzed by bout, average ILI within bouts was not significantly different between sexes for any drinking condition (**Fig.5A**) (MW: AOnly *p=*0.021 n.s. with multiple corrections; AQ10 *p=*0.5949; AQ60 *p=*0.0443; AQ100 *p=*0.7901). However, there was a significant effect across drinking conditions in both sexes (KW, *ps<*0.0001), related to slower licking under higher challenge (see Fig. legend). Similar results were observed when bout speed was examined by session, where average bout ILI was not different for any drinking condition (**Fig.5B**) (MW all *ps>*0.4), but with significant effects across drinking conditions in both sexes (KW, *ps<*0.0001). Thus, even with slightly faster ILI in females when examining ILIs separate from bout (**Fig.4A,B**), females still had sufficient longer ILIs within bouts, and organized their licking within bouts, such that there were no differences in average bout licking speed across the sexes.

**Figure 5.**
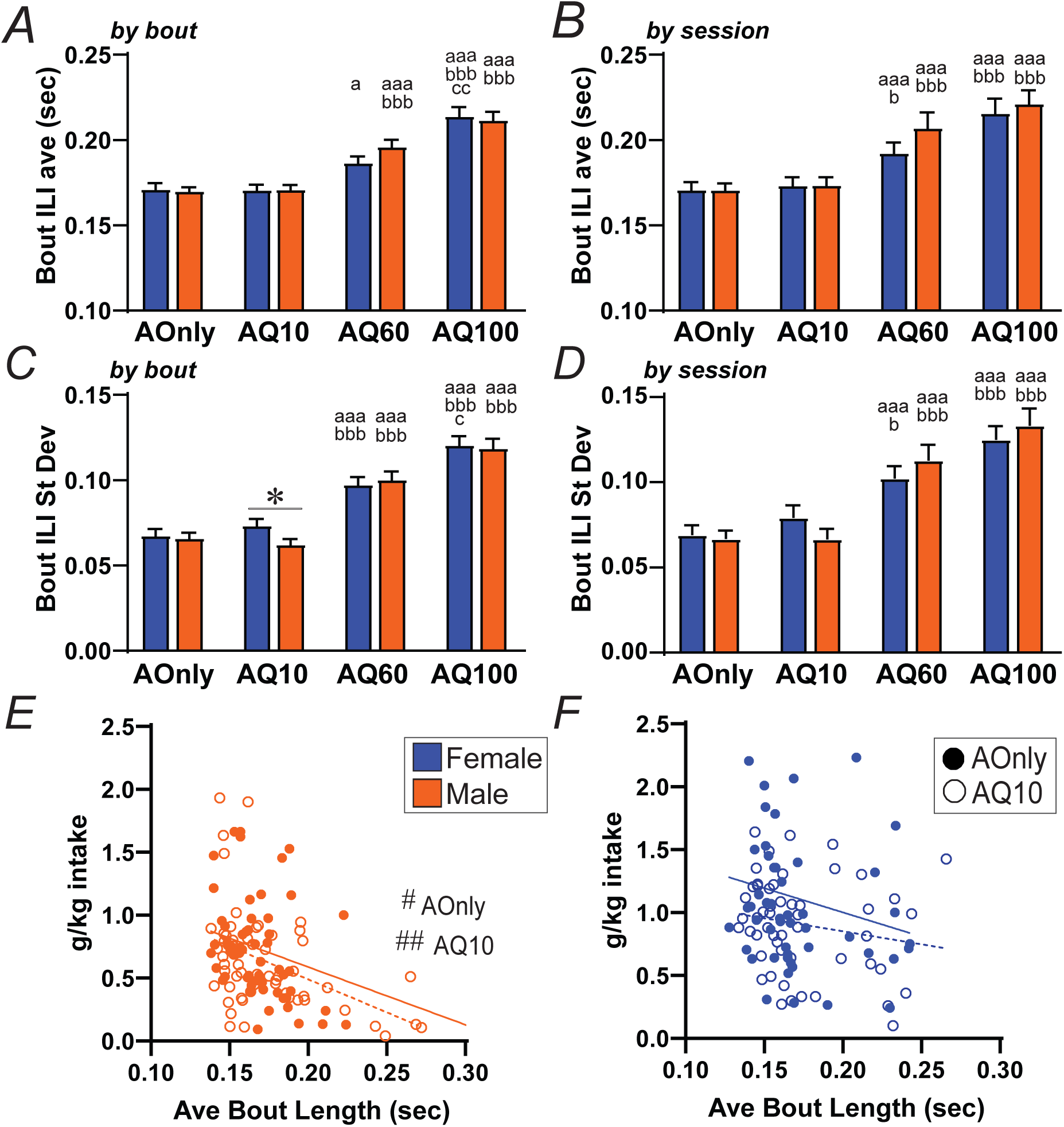
Females are largely similar to males in bout speed and variability when ILIs are averaged within each bout. **(A,C)** Average **(A)** and standard deviation **(C)** of average ILI within each bout, examined by bout. **(B,D)** Average **(B)** and standard deviation **(D)** of average ILI within each bout, examined by session. (E) Slower average bout speed predicted lower intake in males. (F) Average bout speed was unrelated to total intake in females. ∗ *p<*0.0125, MW effect of sex. For letters: a, aaa *p<*0.05, *p<*0.001 effect of drinking condition vs AOD; b, bbb *p<*0.05, *p<*0.001 AQ60 or AQ100 vs AQ10; c, cc *p<*0.05, *p<*0.01 Q100 vs Q60. #,## *p<*0.05, *p<*0.01 correlation.

Variability in licking speed can also be of value as an indicator of motivation [47]. The pattern for ILI standard deviation within bouts was overall similar to that seen for average ILI. When examined by bout, there were no sex differences in ILI variability for AOnly, AQ60, or AQ100 (MW all *ps>*0.2), although there was a significant difference for AQ10 (MW *p=*0.0074) (**Fig.5C**); there were also significant effects across drinking conditions (KW *p<*0.0001 in both sexes). Similarly, when examined by session (**Fig.5D**), average bout ILI variability was not different between females and males for any drinking condition (MW all *ps>*0.16), although there were significant effects across drinking condition in each sex (KW *ps<*0.0001). Thus, there were few significant sex differences in average bout licking speed or its variability.

Since females have longer bouts under AOnly and AQ10 (but not higher challenge) and maintained lick volume under higher challenge (but not AOnly and AQ10), these very limited differences in licking speed are of particular interest. Differences in licking speed have been widely linked to altered motivation, and one interpretation is that females have several forms of greater persistence-like responding for alcohol that does not involve differences in vigor per se. In addition, slightly faster overall licking might indicate greater motivation in females, but, with strong differences in g/kg-per-lick (**Fig.3F**) and body size, the basic geometry and execution of licking may differ in females and males, separate from motivation. Thus, we next examined whether there were correlations between average bout speed and total intake across sessions: decreased intake level with slower responding might suggest a relation between licking rate and some aspect of motivation. Interestingly, for males, the g/kg level was significantly and negatively correlated with average bout speed with AOnly (AOnly: *p=*0.0116, R^2^=0.1085) and AQ10 (AQ10: *p=*0.0007, R^2^=0.1903) (**Fig.5E**), indicating that slower licking did relate to decreased drinking in males. In contrast, g/kg level was not correlated with average bout speed in females for AOnly (*p=*0.1896, R^2^=0.0349) or AQ10 (*p=*0.2003, R^2^=0.0326) (**Fig.5F**). Under higher challenge, male drinking level remained negatively related to average bout speed (AQ60: *p=*0.0006, R^2^=0.1862; AQ100: *p=*0.0037, R^2^=0.1435), while female drinking was related to bout speed for AQ60 (*p=*0.0065, R^2^=0.1443) but not AQ100 (*p=*0.1370, R^2^=0.0412) (**Suppl. Fig.4**). These perhaps surprising results suggested that the action strategy females employed to execute longer bouts under AOnly and AQ10 was largely unlinked from licking speed, further evidence for a very different drinking strategy in females than males, with greater persistence unrelated to vigor in females.

### Sex differences in other licking measures

It would be of interest if other measures showed sex differences, with particular importance if a given measure showed sex differences only under AOnly and AQ10 conditions, or only under AQ60 or AQ100. Such findings might give additional insights into our proposed persistence-related action strategies utilized by females during different alcohol drinking conditions.

Greater drive to lick might be reflected in starting bouts earlier within a session (see Discussion). The timing of the first bout start in a session did not differ between sexes (MW *ps>*0.16) or across drinking conditions (KW female *p=*0.4923, male *p=*0.0693) (**Fig.6A**). In contrast, the average bout start time (examined by bout) was significantly earlier in females versus males for AOnly (MW *p=*0.0008), although not AQ10 (trend, *p=*0.0372), AQ60 (*p=*0.0914) or AQ100 (*p=*0.4098) (**Fig.6B**). There were also changes across drinking conditions in females (KW *p=*0.0041) and males (*p<*0.0001). A similar pattern was seen with the number of stray licks (licks outside of bouts), which was smaller in females for AOnly (MW *p=*0.0029) but not any other drinking condition (all *ps>*0.2) (**Fig.6C**), and with differences across conditions (KW female *p<*0.0001, male *p=*0.0015) including more stray licks under higher challenge (similar to [47]). Together these findings suggest more efficient organization of bout initiation in females under AOnly, while males were slower to initiate in the absence of challenge (as we observed in [47, 55]).

**Figure 6.**
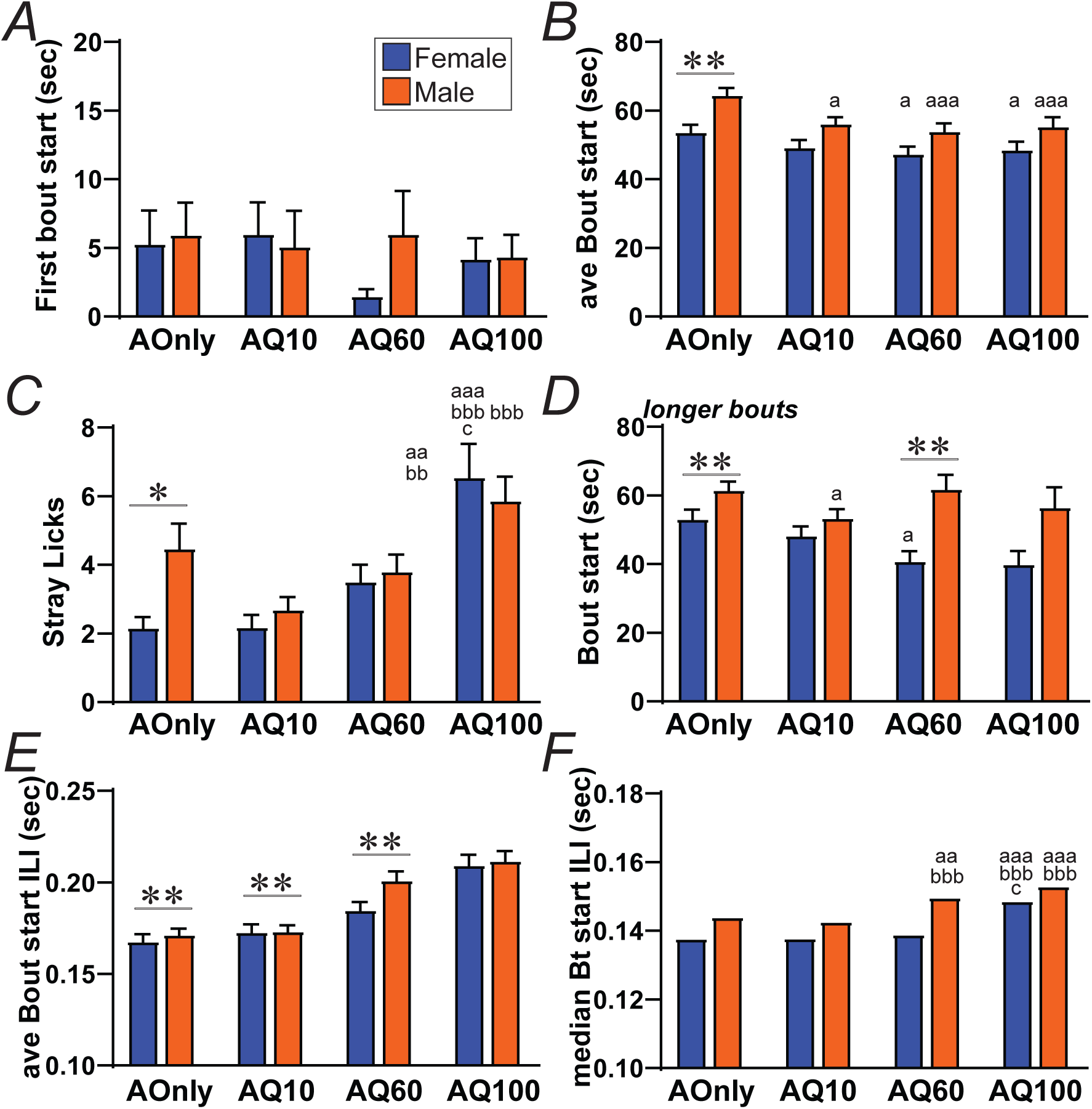
Sex differences in bout start time and speed. **(A)** No difference in time of first bout but **(B)** sooner bouts in females under AOnly. **(C)** Fewer stray licks (licks outside bouts) in females under AOnly. **(D)** Longer bouts (see Methods and Suppl.Figs.5,6) started sooner in females under AOnly and AQ60. **(E,F)** Average **(E)** and median **(F)** of ILI at the beginning of bouts (see Methods), with females having significantly faster bout start under all drinking conditions except AQ100. ∗,∗∗ *p<*0.0125, *p<*0.0025, MW effect of sex. For letters: a, aa, aaa *p<*0.05, *p<*0.01, *p<*0.001 effect of drinking condition vs AOD; bb, bbb *p<*0.01, *p<*0.001 AQ60 or AQ100 vs AQ10; c *p<*0.05 Q100 vs Q60.

We have also extensively examined the properties of bouts of different lengths, divided into shorter bouts (3-9 licks), medium bouts (10-21 licks) and longer bouts (22 and larger bouts) (see [47, 55]). While there were no sex differences in the number of these bout length groups, almost all significant sex differences were seen with longer bouts (**Suppl.Figs.5,6**). In particular, longer bouts started significantly sooner in females versus for AQ60 (MW *p=*0.0004), with a trend for AQ100 (*p=*0.0975); however, longer bouts also started significantly earlier for AOnly (*p=0*.0074), although not AQ10 (*p=*0.3280) (**Fig.6D**) (and KW across conditions female *p=*0.0124, male *p=*0.0210). Thus, females initiated longer bouts significantly earlier under AQ60, with a larger difference between females and males (∼35%) relative to AOnly (∼14%). Further, since measures related to longer and shorter bouts show can correlations with total intake (below), these may suggest other aspects of different female action strategies, including under higher challenge.

Finally, our previous studies have linked faster licking speed at the onset of a bout to greater bout duration [47, 55]. Interestingly, and unlike other measures of lick rate (described above), licking speeds at bout onset (average of the first two ILIs) were slightly but significantly faster in females versus males for AOnly (MW *p<*0.0001), AQ10 (*p=*0.0043), and AQ60 (*p<*0.0001), but not AQ100 (*p=*0.4526) (**Fig.6E,F**), and with significant slowing with higher challenge (KW *p<*0.0001 for both sexes). This may suggest that, while licking speed overall was unlinked from level of responding under AOnly and AQ10, females may in addition have had a particular focus on bout initiation, reflected both in faster bout onset and earlier onset of bouts (above).

### Larger scale relationships among session measures

With the large number of response measures within each session, we can perform larger correlations to indicate whether particular aspects of responding might be more related to higher drinking levels. Thus, we first examined the correlation between g/kg level and other session measures, within each sex/drinking condition, with findings detailed in **Suppl. Tables 2 and 3**. Although the stringency for being considered significant became very high when correcting for multiple comparisons, some interesting patterns were evident. First, total licks were, across the board, highly significantly related to intake level. Second, under higher challenge, greater intake level was related, in both females and males, to having longer bouts, more bouts, and more bouts in the “longer” category; overall, these measures were less related to intake level for AOnly and AQ10. Third, level of alcohol intake was overall significantly related to measures of licking speed and lick speed variability in males, while female intake levels were not. This concurs

### Relation of estrous and ovariectomy to alcohol drinking

Given the higher level of intake in female rats, along with the greater alcohol drinking hazards in women (see Introduction), it is of critical importance to determine the relation of female alcohol drinking to estrous cycle. Some evidence has suggested that sex hormones can significantly modulate intake of intoxicants, especially estrogen (as we recently reviewed in Radke et al., 2021). However, several studies find, once an addiction-related state is established, that responding is unrelated to estrous stage [18,70–74] including with alcohol [75–77]. Here, we examined drinking levels across six estrous stages, proestrus, estrus, late estrus, metestrus, early diestrus, and late diestrus, across 45 female rats, identified as described in Methods and **Suppl. Fig.7**. Importantly, vaginal samples were taken just after drinking, to best match drinking level with estrus stage (which we consider particularly critical since the entire cycle occurs across 4-5 days in rat, introducing potential for mis-identifying estrus stage). Nonetheless, while we attempted to determine drinking for each rat for each stage, some stages (especially proestrus) were underrepresented, perhaps because of the short duration of the proestrus.

As shown in Fig.7, we observed no significant differences across estrous stages for AOnly (**Fig.7A**, KW p=0.1882), AQ10 (**Fig.7B**, KW p=0.2423), or AQ60 (**Fig.7C**, KW p=0.0733) (AQ100 was not examined). Number of samples per stage are in each bar of each graph. Thus, our findings indicate that drinking level does not vary across estrus stage for AOnly, or moderate- or higher-challenge alcohol consumption.

**Figure 7.**
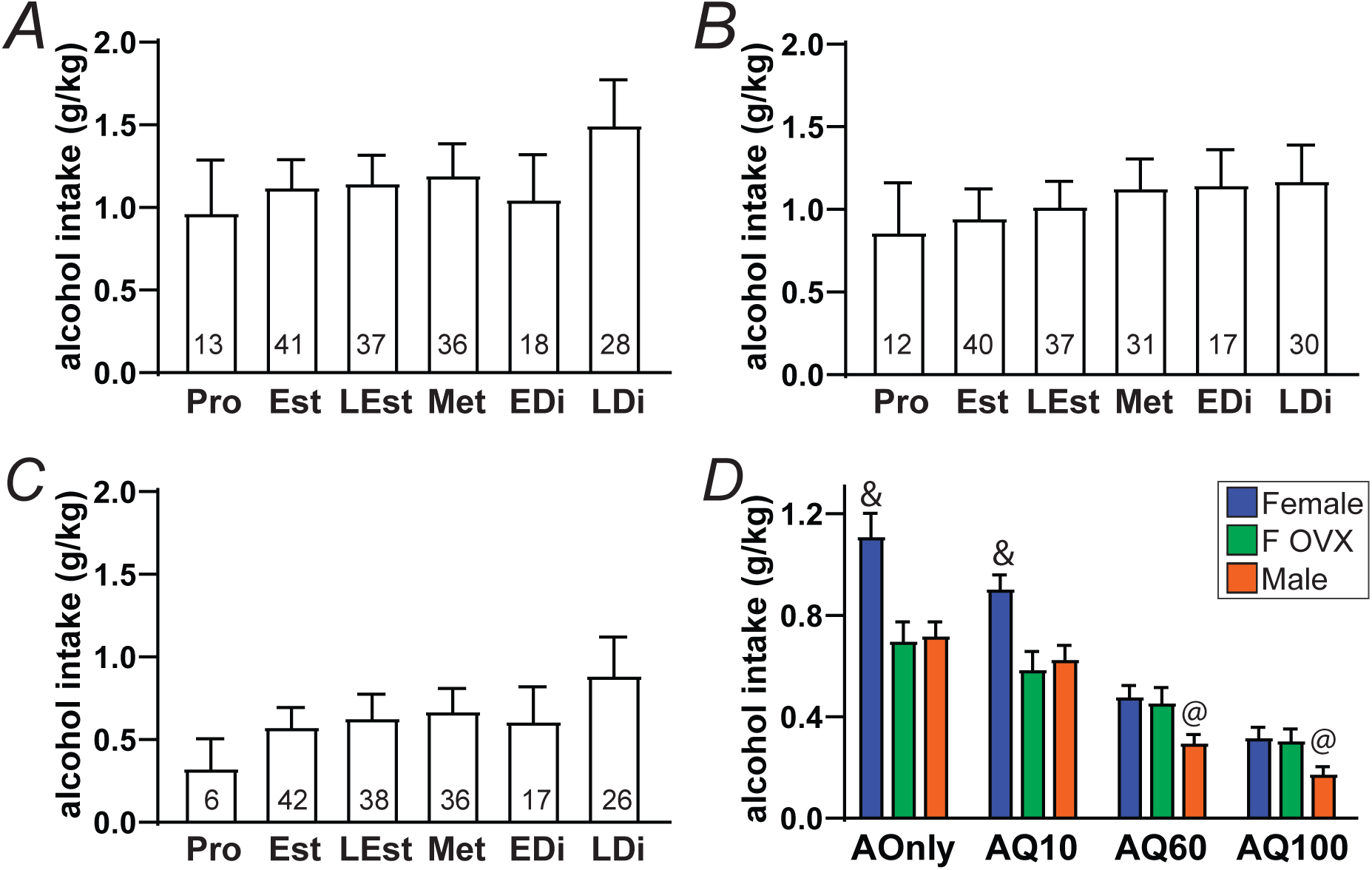
Female drinking in relation to estrous stage and ovariectomy (OVX). **(A-C)** Alcohol drinking levels across six estrus stages for **(A)** AOnly, **(B)** AQ10, and **(C)** AQ60. **(D)** OVX female AOnly and AQ10 intake was significantly reduced relative to intact females, and decreased to male levels, with no OVX effect on AQ60 or AQ100. Male and intact female g/kg intake data are the same as shown in Fig.1B. Pro: proestrus; Est: estrus; Lest: late estrus; Met: metestrus; EDi: early diestrus; LDi: late diestrus. & *p<*0.05 intact female vs OVX or male, @ *p<*0.05 male vs intact or OVX female.

We also examined the impact of ovariectomy (OVX) on consumption (**Fig.7D**, n=14). Interestingly, OVX decreased intake of AOnly and AQ10, such that OVX female g/kg intake was significantly lower than intact females but not different from males under AOnly or AQ10 (KW within each drinking condition *ps<*0.0005; post-hoc *ps<*0.02 intact female vs OVX, *ps>*0.9 OVX vs male,). In contrast, OVX had no impact on AQ60 or AQ100 intake, where OVX females were not different from intact females but significantly greater than males (KW within each drinking condition *ps<*0.002; post-hoc *ps>*0.9 intact female vs OVX, *ps<*0.05 OVX vs male, within each drinking condition). Thus, it is tempting to speculate that some aspect of ovary-dependent signaling plays an important role in the bout lengthening under AOnly and AQ10 in females (see Discussion).

## DISCUSSION

While female humans and rodents have historically been understudied, rates of problem alcohol drinking have been increasing in recent decades, and women can have greater alcohol problems including negative health impacts (see Introduction). Thus, there is considerable necessity in understanding where alcohol consumption involves similar or different cognitive/emotional strategies and underlying mechanisms, which would be of great value when seeking to develop more effective, personalized treatments, since existing treatments for AUD are of limited efficacy. Here, we used large cohorts of adult Wistar rats (28 females, 30 males) and lickometry to robustly assess whether there were sex differences in patterns of responding for alcohol-only and under compulsion-like drinking with moderate or higher challenge. Overall, females had a similar level total licking as males, with higher alcohol intake, for each drinking condition. There were also no sex differences in sensitivity to quinine in alcohol, or quinine in water, and both females and males both showed significantly greater tolerance of quinine in alcohol and dislike of quinine in water. Even with these overall sex similarities, more specific response measures indicated that females and males may have had different action strategies. In particular, females had significantly longer bouts (40-50% longer than males) under alcohol-only and moderate challenge, with no sex differences under higher challenge. Conversely, under higher challenge, females retained several aspects of responding not seen in males, including more retaining higher g/kg intake level per lick, and earlier onset of longer bouts. Furthermore, licking speed has been associated with motivation, and while females overall licked slightly faster, licking speed averaged within-bout showed no sex differences, even though licking speed slowed significantly under challenge in both females in males, suggesting that lick speed can change in relation to the motivational environment. In addition, female intake level under alcohol-only and moderate challenge was unrelated to licking speed, while slower licking in males predicted lower intake. We interpret these patterns as indicating sex differences in underlying action strategy, where females exhibit greater persistence-like responding but not action vigor per se, and, interestingly, with different strategies under lower versus higher challenge. Finally, drinking levels did not differ across the estrous cycle, although ovariectomy reduced alcohol-only and moderate-challenge intake. Thus, there were several aspects of responding for alcohol that were sex-similar, suggesting some common drinking mechanisms, but there was clear, somewhat more nuanced, evidence for sex-different alcohol response strategies, and we propose that these responding differences might make an outsized contribution to excessive drinking since women can have more drinking problems. Thus, our studies provide important context for future work examining sex differences in mechanisms underlying different aspects of pathological alcohol drinking.

Together, these results agree with our previous studies that male rats do not reduce intake level under AQ10, and thus we have called this form of drinking aversion-resistance under moderate challenge [47, 55]. Also, while Q60 and Q100 significantly reduced intake when added to alcohol, water intake was greatly reduced by addition of 10 and 60 mg/L quinine, indicating that AQ60 intake could still be considered aversion-resistant even while intake was somewhat decreased relative to AQ10. Our results agree with previous findings showing no sex differences in basic aversion sensitivity for quinine [78–80] or shock [81]. We also find that aversion-resistance across doses of quinine was not different between females and males, agreeing with previous studies showing sex-similar quinine sensitivity in alcohol under home cage bottle-drinking conditions ([80,82,83] but see [75]). However, other studies have found that females can be more aversion-resistant, including under operant responding for alcohol [79] and shock-resistance in a place preference model [81]. At present, the reason for the mixed results remains unclear. However, one interesting possibility is where female responding for alcohol is more driven by alcohol cues, while male responding is regulated by both alcohol cue and context [84], including where female and male humans can show similar behavior under simpler response requirements, but differences in more complex tasks (discussed in [84]). In this view, female focus on a specific landmark (cue) within a situation may lead to quicker or stronger learning, e.g. in an operant task for alcohol, and this may contribute importantly to greater female aversion-resistance in rodents and level of addiction in humans (although there were no sex differences in quinine reduction of breakpoint for alcohol [76]). Our findings here are thus significant because, even though macroscopic quinine resistance did not differ between female and male rats, we still observed differences in behavioral response patterns that indicate different behavioral strategies in females relative to males.

One more general challenge is whether rodent drinking models reflect mechanisms in human problem intake. The intermittent access alcohol model we use exhibits several features considered to relate to human AUD, including escalating intake [36, 85], sensitivity to compounds that reduce human drinking [85], withdrawal symptoms (although moderate) [86, 87], and front loading (indicating motivation for alcohol [47,55,88,89]). However, CLAD models have been challenged because negative consequences for humans are often in the future, unlike acute in rodent models. However, an alternate perspective is that, for treatment seekers, the negative consequences are more acute [27] (“actively struggling with the balance between known bad outcomes and the desire to drink” [25]), and we have always considered our findings primarily to be more relevant for treatment seekers. Thus, some clinicians have expressed support for the value of rodent CLAD models [31,48,90–92]. In addition, one group found that people with AUD were not more insensitive to cost [42, 43] which could question the importance of consequence-resistance. However, intake despite negative consequences features prominently in the DSM-V definition of AUD (e.g. [37, 38]), and other groups do find less sensitivity to cost with AUD [39–41], while greater drinking is associated with more alcohol problems [38,44–46]. Thus, [31, 92] have emphasized, from a clinician’s perspective, that multiple factors can contribute to AUD expression [93–99] including compulsion-like drives. Also, we utilize quinine-resistant drinking, rather than footshock-resistance, since similar cortical-NAcb projections mediate both quinine- and footshock-resistant alcohol consumption [50], and higher shock-resistance correlates with higher quinine-resistance for alcohol [90, 91], which together validate the technically simple quinine-resistance to study CLAD (which is also easier to test in a graded manner vs shock).

Our previous studies using lickometry to help understand behavioral strategies underlying AOnly and CLAD found that AQ10 responding (in male rats) showed reduced variability in many response measures. This contributed to our development of the “head down and push” model of CLAD, where the rat quickly identifies that it is licking challenge-adulterated alcohol, and changes response strategy to sustain licking in an attempt to get sufficient alcohol while minimizing attention to and impact of negative consequences [47, 55]. However, we should note that these previous studies used a licking apparatus with a more complicated geometry (licking in a Med Associated operant box, where the rat had to rear slightly and insert it’s nose into a space to lick). Nonetheless, there are several consistent patterns with present studies, especially where higher challenge reduced licking speed, overall intake, and led to bout disorganization (the presence of more shorter bouts and fewer longer bouts under higher challenge).

The observation of different behavioral response strategies in females versus males also suggests potential sex differences in brain circuits that promote alcohol drinking. Many recent studies have examined brain mechanisms for female and male alcohol drinking, where some pathways play a similar role across the sexes, while a number of sex differences have been observed (reviewed in [18,25,100–103], e.g. for norepinephrine [104] and orexin [25, 105] which can be important regulators of alcohol drinking. It is also interesting that human CLAD studies implicate insula circuitry in both women and men [48, 49], agreeing with insula importance for CLAD in rodents [50,52,53,59], and we [59] and others [106] also find insula importance for promoting AOnly, consistent with the importance of the insula-related salience network in motivated behavior more generally [54,107–110]. Nonetheless, there are a wide range of brain regions and signaling pathways that contribute to alcohol intake. Thus, when taken together, studies of sex-related responding indicate there are important and diverse differences in the cognitive/emotional strategies that females and males utilize to promote different aspects of problem alcohol consumption, which warrants considerable additional attention to uncover the specific mechanisms that underly such sex differences.

We also find no influence of estrous stage on AOnly or CLAD. While estrous cycle can influence alcohol drinking in some studies (reviewed in [25]), our findings are consistent with other studies finding that estrous cycle can have limited influence on rodent addiction-related behaviors once established [18,70–74] including alcohol [75–77]. However, we found that ovariectomy did significantly reduced alcohol drinking only under AOnly andAQ10, reducing the longer bouts to male levels, with no effect on intake under higher challenge. Thus, we could speculate that some aspect of ovary-dependent signaling contribute importantly to the bout lengthening observed under AOnly and AQ10 in females. However, the observation of no differences in drinking across the estrous cycle suggests that any such influences are unlikely due simply to e.g. higher estrogen or progesterone (which would be predicted to impact intake differently across the cycle). Indeed, other studies have found no differences in alcohol drinking across the cycle in intact females, but reduced drinking after OVX (which increased to intact levels when estrogen was restored) [77]. Thus, such findings important clues about the importance of ovarian hormones for alcohol drinking, while the exact underlying mechanisms remain uncertain. Problem alcohol consumption is a major social harm, and it is critical to understand possible sex differences in the cognitive/emotional strategies that promote alcohol intake, since this may indicate different brain mechanisms which would require selective therapeutics to best address. By comparing a large sample of female and male rats, and the microstructure of licking, we find evidence that females utilize different action strategies than males for AOnly/AQ10 and higher challenge, both suggesting greater persistence-like behavior, including much longer bouts under AOnly/AQ10 which is lost under higher challenge, but adjustments in intake per lick and bout initiation under higher challenge. Interestingly, neither change in action strategy involves changes in vigor per se (as assessed by licking speed). Also, female intake was similar across the estrous cycle. Together, our studies provide important context for future work examining sex differences in pathological drinking mechanisms.

## ACKNOWLEDGEMENTS

We thank Jacob Kellner for assistance with vaginal cytology.

## AUTHOR CONTRIBUTIONS

TDOS and FWH designed the experiments; TDOS, VdPS, MdCA, DM and SW performed the experiments; TDOS, DD, VdPS, MdCA, DM, SW and FWH analyzed the data; TDOS and FWH wrote a first draft of the manuscript, all authors edited the manuscript and approved the final version.

## Suppl. Figure Legends

**Figure S1.**
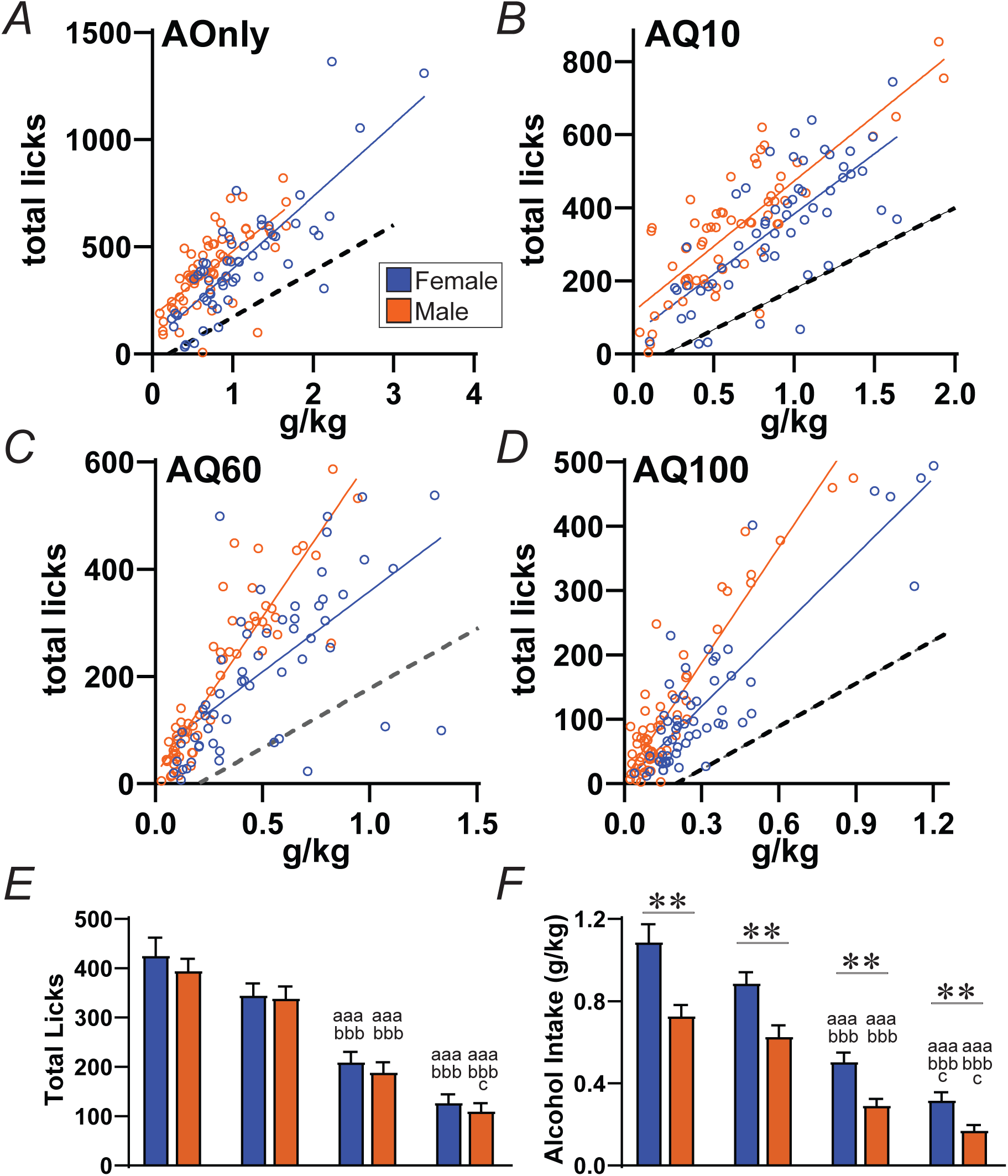
Relation between alcohol intake (g/kg) and total licks, showing high-intake/low-licking sessions that were excluded from analyses for licking parameters. A very small number of sessions showed quite high g/kg relative to total licks, which could suggest the tongue stayed in contact with the sipper tube (and not registering separate licks). There were also sessions with very few licks (<10) which had no bout. All such sessions fell to the right of the dotted lines shown. The number of such excluded sessions was 5, 4, 6, 1 for AOnly, AQ10, AQ60, and AQ100 for females, and 2, 3, 1, 3 for males (leaving >50 for each condition). **(A-D)** The slope relationship for intake and total licks was not different between sexes for **(A)** AOnly or **(B)** AQ10 (*p>*0.1), but did differ significantly between sexes for **(C)** AQ60 and **(D)** AQ100 (*ps<*0.001). The overall patterns for **(E)** total intake and **(F)** alcohol intake, when including all sessions, were similar to those shown in **Fig.1A,B**. ∗∗ *p<*0.0025 MW effect of sex. For letters: a, aa, aaa *p<*0.05, *p<*0.01, *p<*0.001 effect of drinking condition vs AOD (KruskalWallis with Dunn’s post-hoc); b, bb, bbb *p<*0.05, *p<*0.01, *p<*0.001 AQ60 or AQ100 vs AQ10; c, cc *p<*0.05, *p<*0.01 Q100 vs Q60.

**Figure S2.**
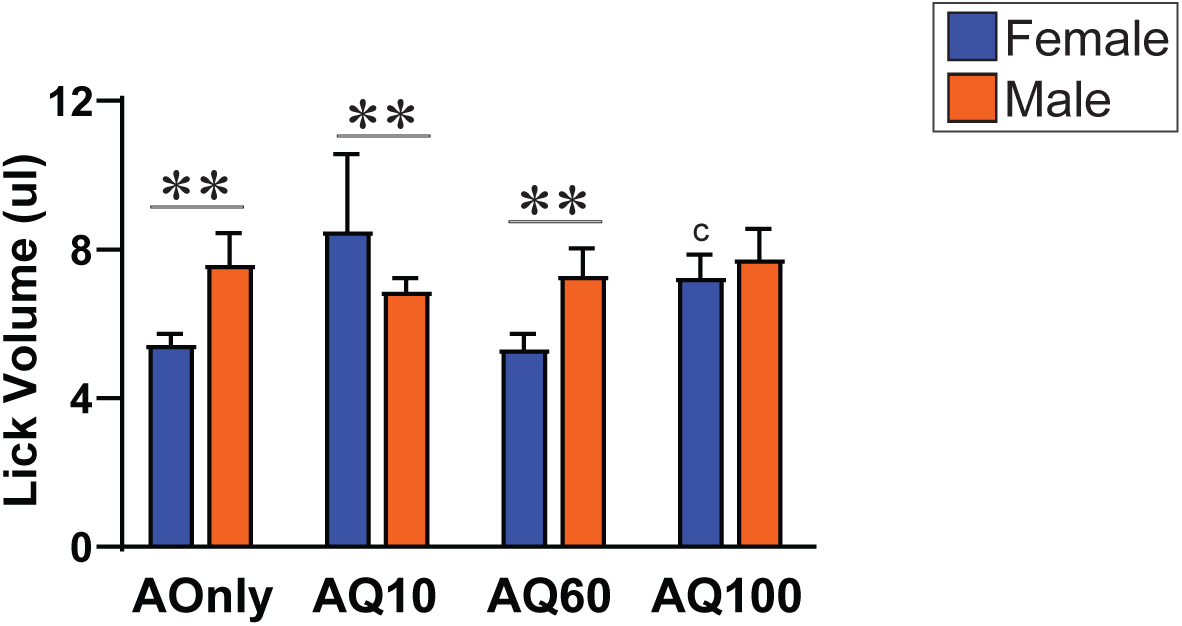
**Lick Volume (volume alcohol consumed/total licks)** measures were somewhat more complex, with females having significantly lower lick volume for AOnly (*p=*0.0008) and AQ60 (*p=*0.0004), males having greater for AQ10 (*p=*0.0010), and no differences for AQ100 (*p=*0.9296), and lick volume was not different across drinking conditions in either sex (KW female *p=*0.0198 with no signif post-hocs, male *p=*0.1986). Lick volume measures may be complicated by differing body sizes in females and males, and thus our main analysis focused on g/kg-per-lick, since this accounts for smaller body size in females. See Fig.S1 for statistical symbols.

**Figure S3.**
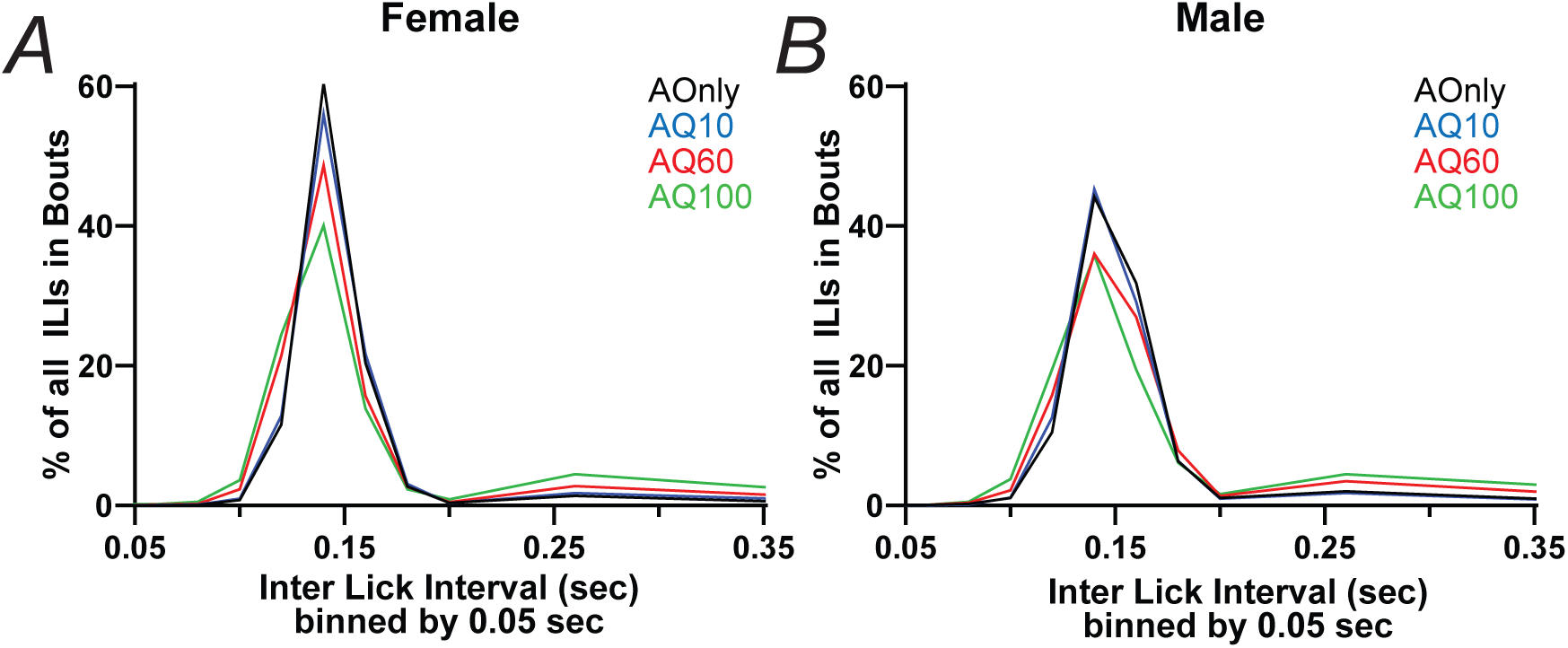
**Histogram of ILI values within bouts**, taking each ILI separately from bout and session. Higher challenge led to decreasing number of ILIs in the main peak at ∼150msec, and increasing ILIs longer than 200 msec. Distribution extends to 999msec, but is truncated at 350 msec.

**Figure S4.**
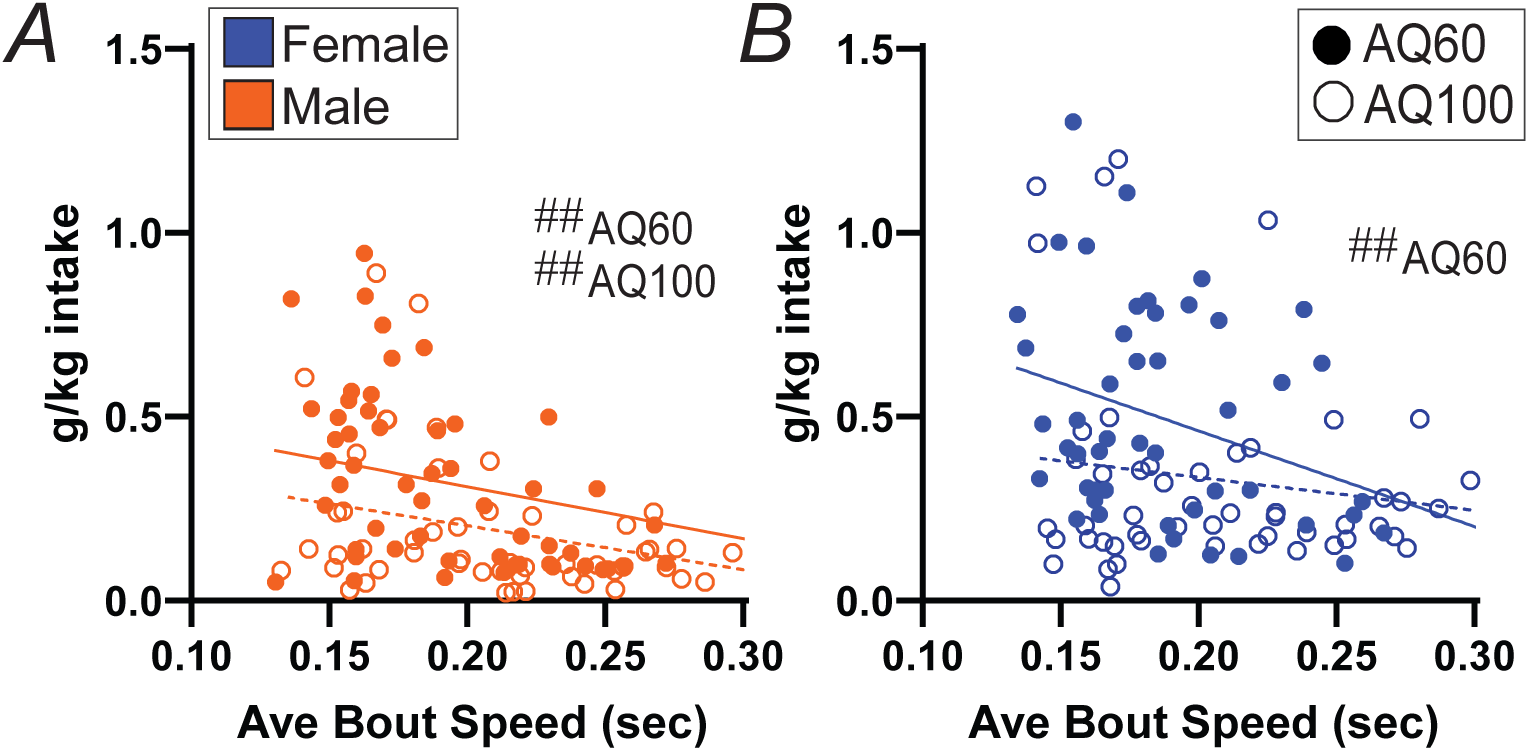
Relation between intake level and average licking speed within bouts under higher challenge conditions. **(A)** In males, average bout speed was negatively correlated with intake level for both AQ60 and AQ100. **(B)** In females, average bout speed was negatively correlated with intake level for AQ60, but not related to intake for AQ100. ## *p<*0.001 for correlation.

**Figure S5.**
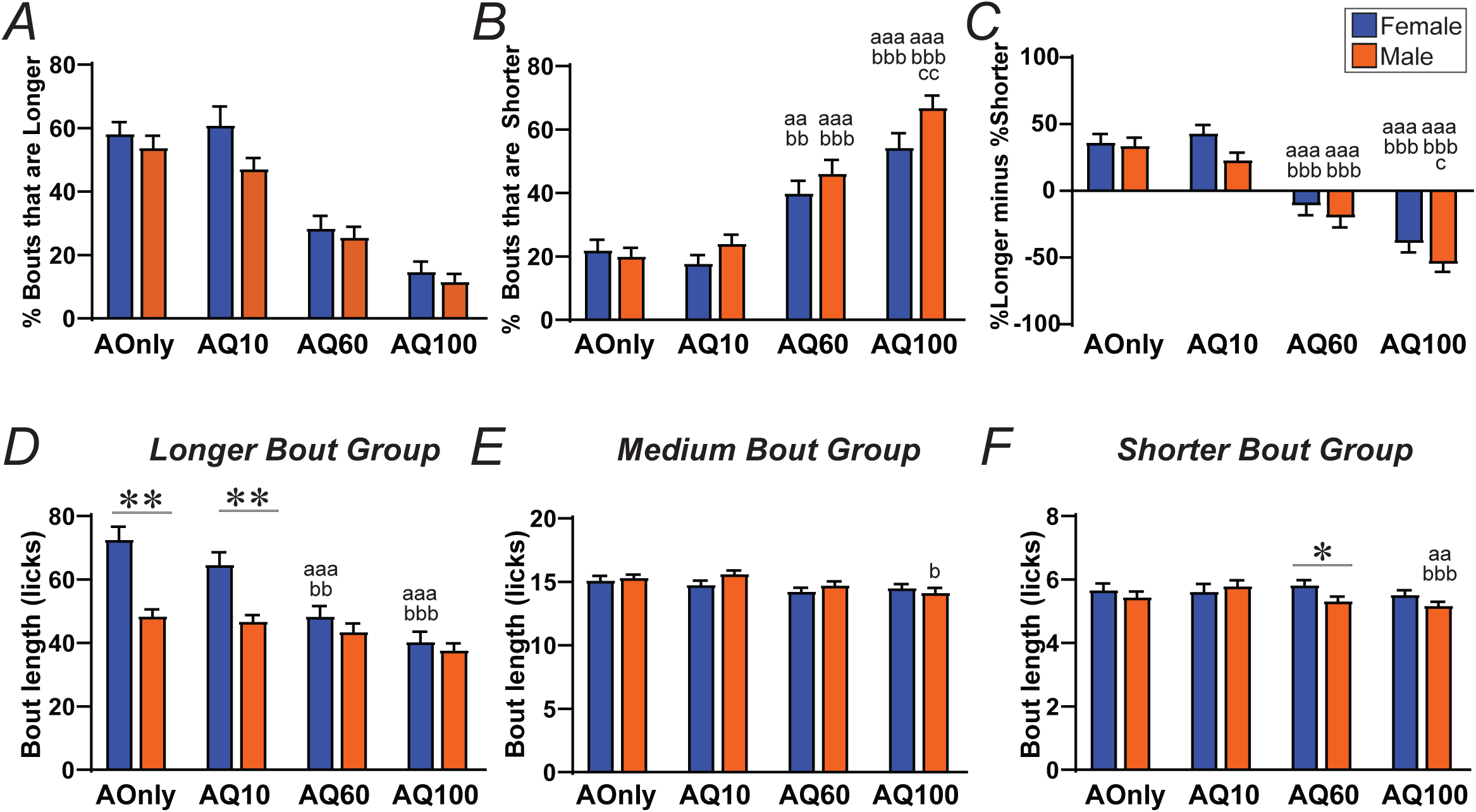
**Dividing bouts into those that are longer (22+ licks), medium (10-21 licks), or shorter (3-9 licks),** as in Darevsky et al., 2019, Darevsky and Hopf, 2020. As observed before, higher challenge conditions have **(A)** fewer longer bouts (% of all bouts in that session) and **(B)** more shorter bouts (% of all bouts in the session), and with **(C)** showing %longer minus %shorter bouts in the session. These patterns, which we have interpreted as evidence of bout fragmentation (Darevsky et al., 2019; Darevsky and Hopf, 2020), were not different between sexes, although with significant differences across drinking conditions (KW *ps<*0.0001). **(D-E)** Average length of bouts within each of the three bout length groups, **(D)** Longer, **(E)** Medium, and **(F)** Shorter. The primary difference was **(D)** lengthier Longer bouts in females for AOnly and AQ10 (as in Fig.2), while there was a small but significant difference in (**F)**. See Fig.S1 for statistical symbols.

**Figure S6.**
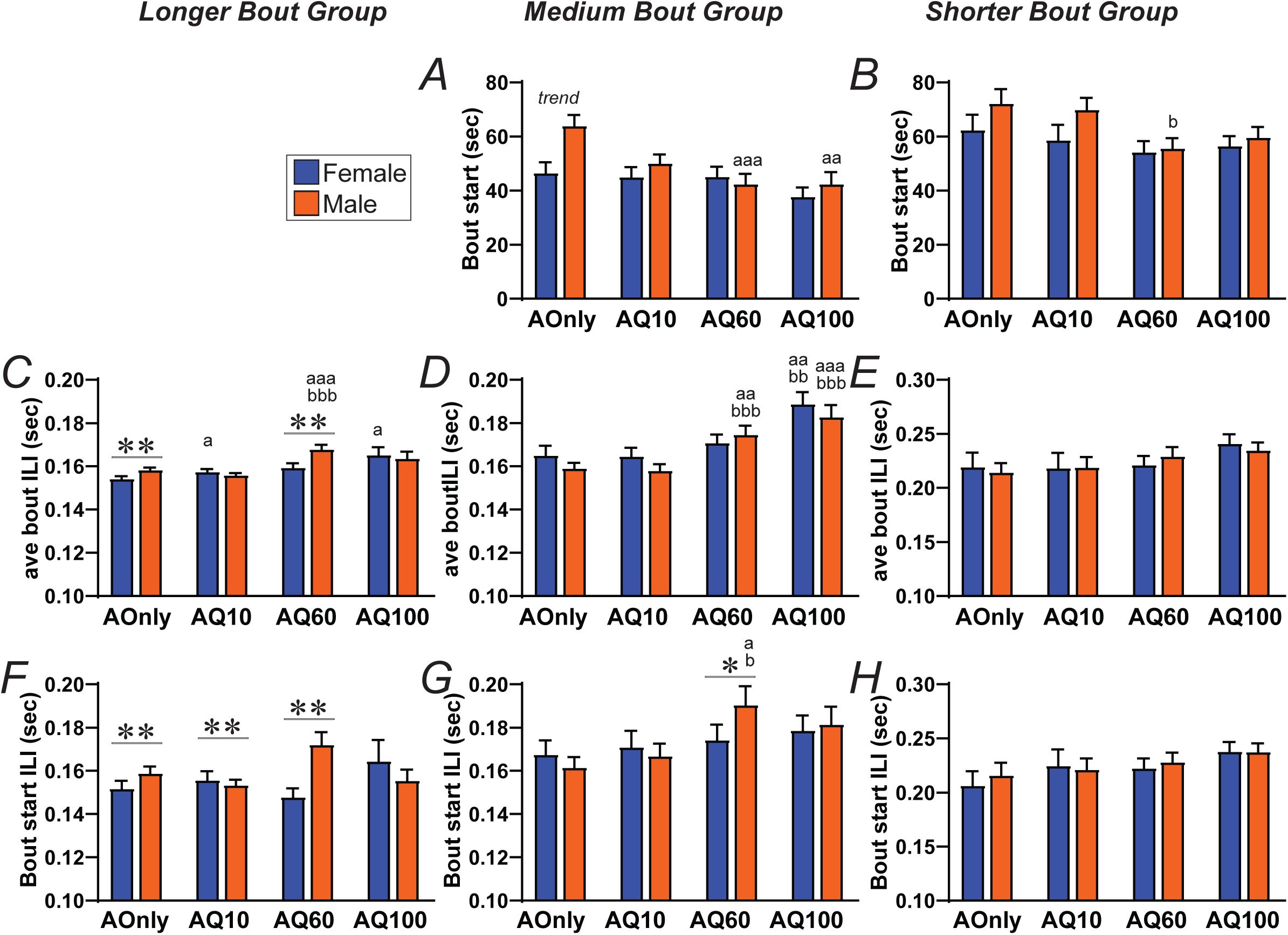
Other properties of bouts that are longer (22+ licks), medium (10-21 licks), or shorter (3-9 licks). **(A,B)** Average bout start time for **(A)** medium bouts (KW across drinking conditions: female *p=*0.2146, male *p<*0.0001) and **(B)** shorter bouts (KW female *p=*0.6573, male *p=*0.0078). **(C-E)** Average bout speed for **(C)** longer bouts (KW female *p=*0.0032, male *p<*0.0001), **(D)** medium bouts (KW female *p=*0.0009, male *p<*0.0001), and **(E)** shorter bouts (KW female *p=*0.2082, male *p=*0.5471). **(F-H)** Average bout speed for **(F)** longer bouts (KW female *p=*0.0709, male *p=*0.0515), **(G)** medium bouts (KW female *p=*0.6509, male *p=*0.0026), and **(H)** shorter bouts (KW female *p=*0.1757, male *p=*0.5238). See Fig.S1 for statistical symbols.

**Figure S7.**
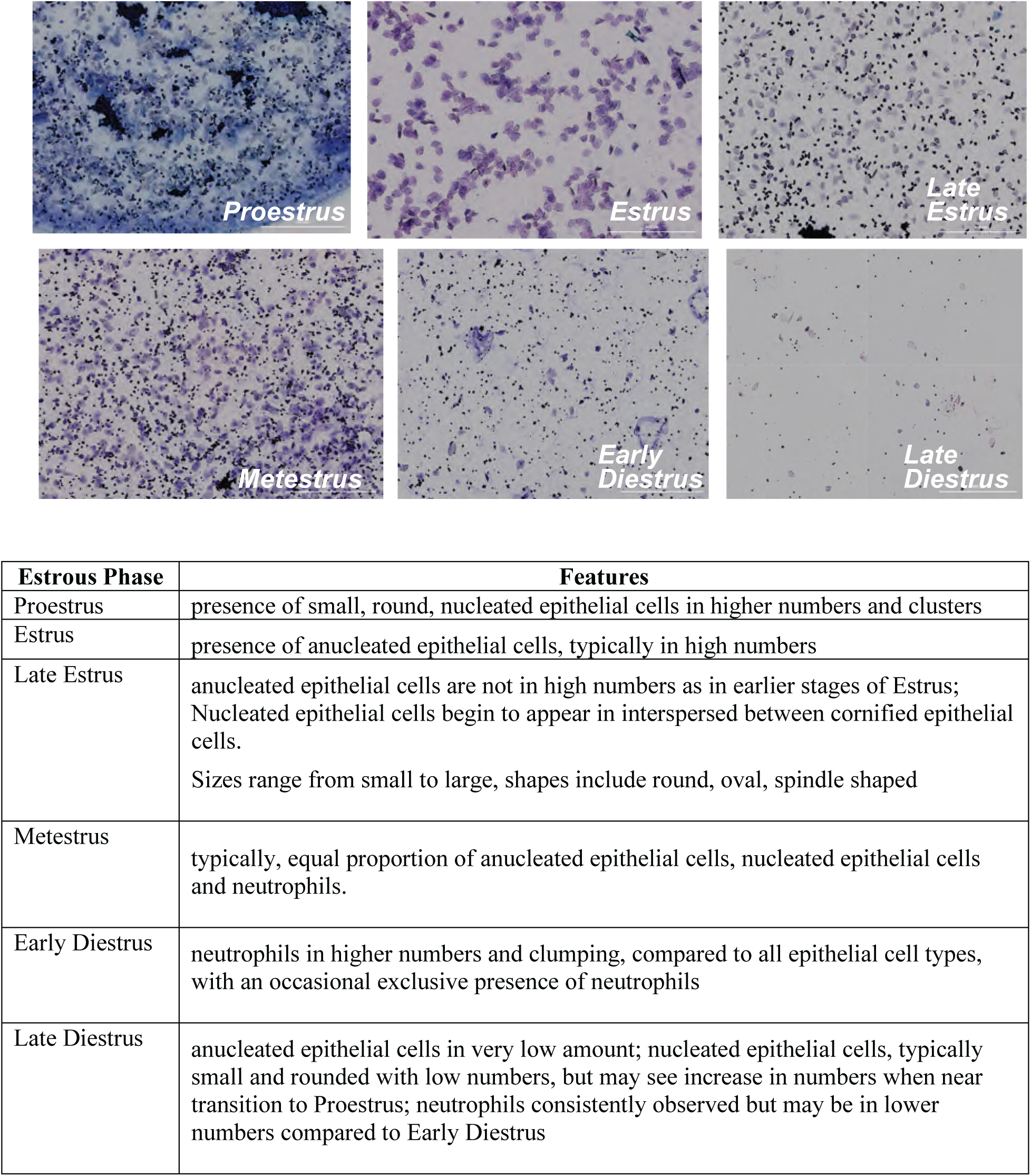
Criterion used to assess different estrus stages. **(Top)** Example images of different stages of the estrus cycle. **(Bottom)** Table summarizing critical factors to distinguish different estrus stages. Briefly, proestrus was defined by the presence of small, round, nucleated epithelial cells. Estrus by the presence of anucleated epithelial cells, and late estrus by the presence of anucleated cells with sizes ranging from small to large and shapes including round, oval and spindle. Metestrus was characterized by the presence of anucleated, epithelial cells and neutrophiles in the same proportion. Early diestrus was defined by the presence of neutrophils, typically in higher numbers and clumping, compared to all epithelial cell types. Late diestrus defined by the presence of lower number of anucleated epithelial cells and neutrophils compared to early diestrus. We note that estrus stage was determined separately by two researchers who were blind to the drinking condition

**Suppl. Table 1.** Differences across IN and CA cohorts. We note that there were significant overall effects across IN and CA cohorts. In this table, we compared differences between the entire IN vs entire CA data set within each sex (i.e., collapsing across drinking condition). We then assessed whether there were differences across the IN and CA data within each sex. With multiple corrections (considering the same 8 variables from the PCA, and 4 conditions), p<0.00156 is needed to be considered significant. Bold shows significant, nonbold nonitalics is trend, and italics is non significant. More generally, for the first 6-7 measures, differences between IN and CA cohorts were similar across sex, especially with greater total g/kg intake and total licks in IN, without differences in lick volume measures or average bout start time. In Suppl. Table W, when examining the relationship between total intake and other session measures (collapsed across sex and drinking condition), lick volume and average bout start time were not significantly related to g/kg, while all other measures were highly related. Further, when examining g/kg versus other session measures within each sex/drinking condition separately, total licks was across the board highly significantly related to intake. Thus, since the largest % differences across cohorts in this table were in g/kg and total licks, it may be that greater intake overall in IN was the primary factor underlying changes in other measures. Further, this table shows average values for each session measure in IN and CA, and, even with differences across cohorts, macroscopic differences between males and females were retained. For example, in Suppl. Table W, greater intake in males related to faster licking, while female intake did not, and, in this Table, males showed differences in lick speed measures, while females did not. This may support the possibility that female licking was more disconnected from licking rate, while male intake level was impacted by licking speed, suggesting different action strategies across the sexes. Thus, taken together, while there are certainly limitations to comparing across the cohorts, the large sample sizes and overall sex-related response differences still provide valuable evidence for potential sex differences in response strategy during alcohol consumption.

|  | FEMALE<br>p value | ave<br>IN | ave<br>CA | %<br>Chng | MALE<br>p value | ave<br>IN | ave<br>CA | %<br>Chng |
| --- | --- | --- | --- | --- | --- | --- | --- | --- |
| Ave Bt Start Time | *0.0921* | 5.0 | 3.6 | 37 | *0.6605* | 4.7 | 6.0 | -22 |
| Lick Vol | 0.0057 | 6.3 | 7.1 | -11 | *0.5916* | 6.9 | 8.1 | -15 |
| Gkg per Lick | *0.2389* | 2.7 | 2.7 | -3 | 0.0428 | 1.8 | 1.9 | -8 |
| Total Lick | **<0.0001** | 361.7 | 228.7 | 58 | **<0.0001** | 315.4 | 218.7 | 44 |
| G/kg | **<0.0001** | 0.916 | 0.497 | 84 | **<0.0001** | 0.593 | 0.322 | 84 |
| # Bouts | 0.0017 | 10.9 | 8.8 | 24 | 0.0083 | 12.3 | 10.1 | 21 |
| Longest Bt | 0.0024 | 106.6 | 78.5 | 36 | **0.001** | 67.2 | 58.3 | 15 |
| Ave Bt Length | 0.0031 | 37.0 | 30.7 | 20 | **<0.0001** | 26.5 | 21.5 | 23 |
| Ave Bt Speed | 0.022 | 0.181 | 0.196 | -8 | **<0.0001** | 0.170 | 0.216 | -21 |
| Ave Bt Stdev | *0.5262* | 0.091 | 0.099 | -8 | **<0.0001** | 0.070 | 0.120 | -41 |
| Weight | **<0.0001** | 314.7 | 411.5 | -24 | **<0.0001** | 581.1 | 684.0 | -15 |
| % Bts Long | 0.0325 | 43.83 | 36.56 | 20 | **<0.0001** | 42.1 | 27.3 | 54 |
| %Long - %Short | 0.0054 | -17.6 | 5.1 | -446 | **<0.0001** | -13.55 | 22.888 | -159 |
| % Bts Short | **0.0003** | 26.2 | 41.7 | -37 | **<0.0001** | 28.5 | 50.2 | -43 |
| Ave Bt Start Speed | 0.0037 | 0.176 | 0.200 | -12 | **<0.0001** | 0.173 | 0.219 | -21 |
| Ave Bt Var | *0.6267* | 0.018 | 0.020 | -11 | **<0.0001** | 0.013 | 0.028 | -53 |
| Stray Licks | 0.0123 | 2.85 | 4.47 | -36 | **<0.0001** | 2.7 | 5.8 | -54 |

**Suppl. Table 2.** Alcohol intake (g/kg) correlated with other session measures. Since we consider total drinking level (in g/kg alcohol consumed) of primary importance, we performed a series of correlations to determine whether g/kg was significantly related to other session-level measures. We did this two ways. First, we collapsed across sex and drinking condition, and the left two data columns show the p value and slope for g/kg versus each session measures. Lick volume and average bout start time were not significantly related to g/kg, while all other measures were highly related. Second, we examined the g/kg correlation with each session measure separately for each sex/drinking condition. **Suppl. Table 3** shows the actual p values for each condition, here we summarize the data, condensing across relatively lower challenge conditions (AOnly+AQ10) vs higher challenge conditions (A+AQ10). “F” or “M” in the right columns means g/kg was significantly correlated with the session measure. “(M)” means one of two drinking conditions was significant in males, with the other showing a strong trend [similar for “(F)”]. While the very high stringency for significance (p<0.00078, see **Suppl. Table 3**) does complicate interpretation, there are some clear patterns. First, total licks was across the board highly significantly related to intake. Second, under higher challenge, greater intake was related, in both females and males, to having longer bouts, more bouts, and more bouts in the “Longer” category; overall, these measures were less related to intake level for AOnly and AQ10. Third, in males, level of alcohol intake was overall significantly related to measures of licking speed and lick speed variability, while female intake levels were not.

|  | All Sessions |  |  | signif p value when analyzed separated by sex & drink cond |  |
| --- | --- | --- | --- | --- | --- |
|  | p value | slope | more intake assoc with | AOnly+AQ10 | AQ60+AQ100 |
| Ave Bt Start Time | 0.8838 | -0.243 | NOT bout start |  |  |
| Lick Vol | 0.8984 | -0.085 | NOT lick volume |  |  |
| Gkg per Lick | 0.0803 | 0.2094 | NOT lick volume | (M) |  |
| Total Lick | <0.0001 | 375.2 | more licks | F M | F M |
| # Bouts | <0.0001 | 4.2 | more bouts | (M) | F M |
| Longest Bt | <0.0001 | 89.2 | longer bouts |  | F M |
| Ave Bt Length | <0.0001 | 29.9 | longer bouts |  | F M |
| Ave Bt Speed | <0.0001 | -0.044 | faster bouts | (M) | (M) |
| Ave Bt Stdev | <0.0001 | -0.058 | less variable bouts | M | (M) |
| Weight | <0.0001 | -127.5 | weighing less | (F) M |  |
| % Bts Long | <0.0001 | 38.5 | more long bouts |  | F M |
| %Long - %Short | <0.0001 | -75.2 | more long bouts | M | F M |
| % Bts Short | <0.0001 | -36.7 | more long bouts | M | F M |
| Ave Bt Start Speed | <0.0001 | -0.048 | faster bouts |  |  |
| Ave Bt Var | <0.0001 | -0.018 | less variable bouts |  | (M) |
| Stray Licks | <0.0001 | -2.4 | less stray licks |  |  |

**Suppl. Table 3.**
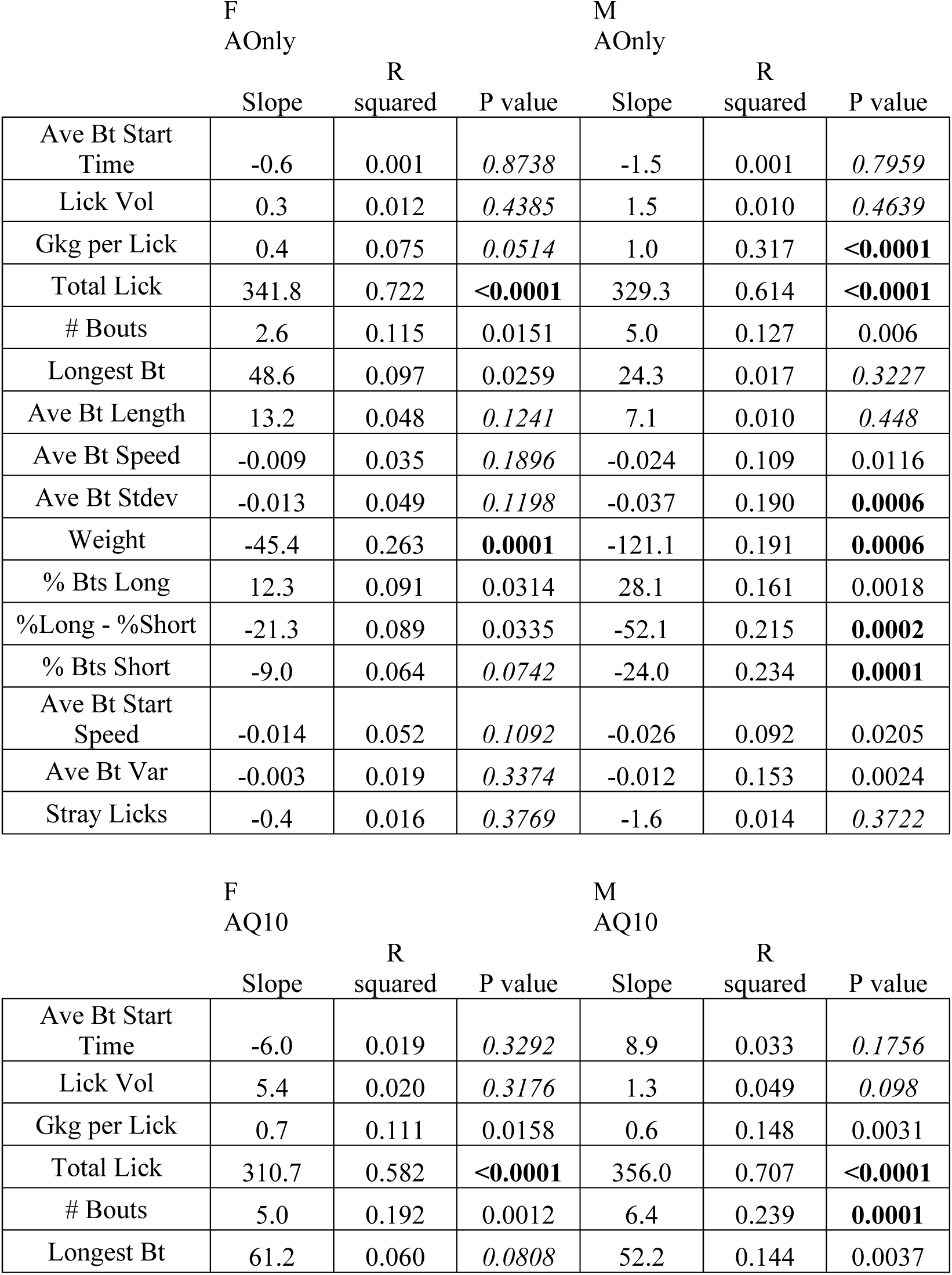
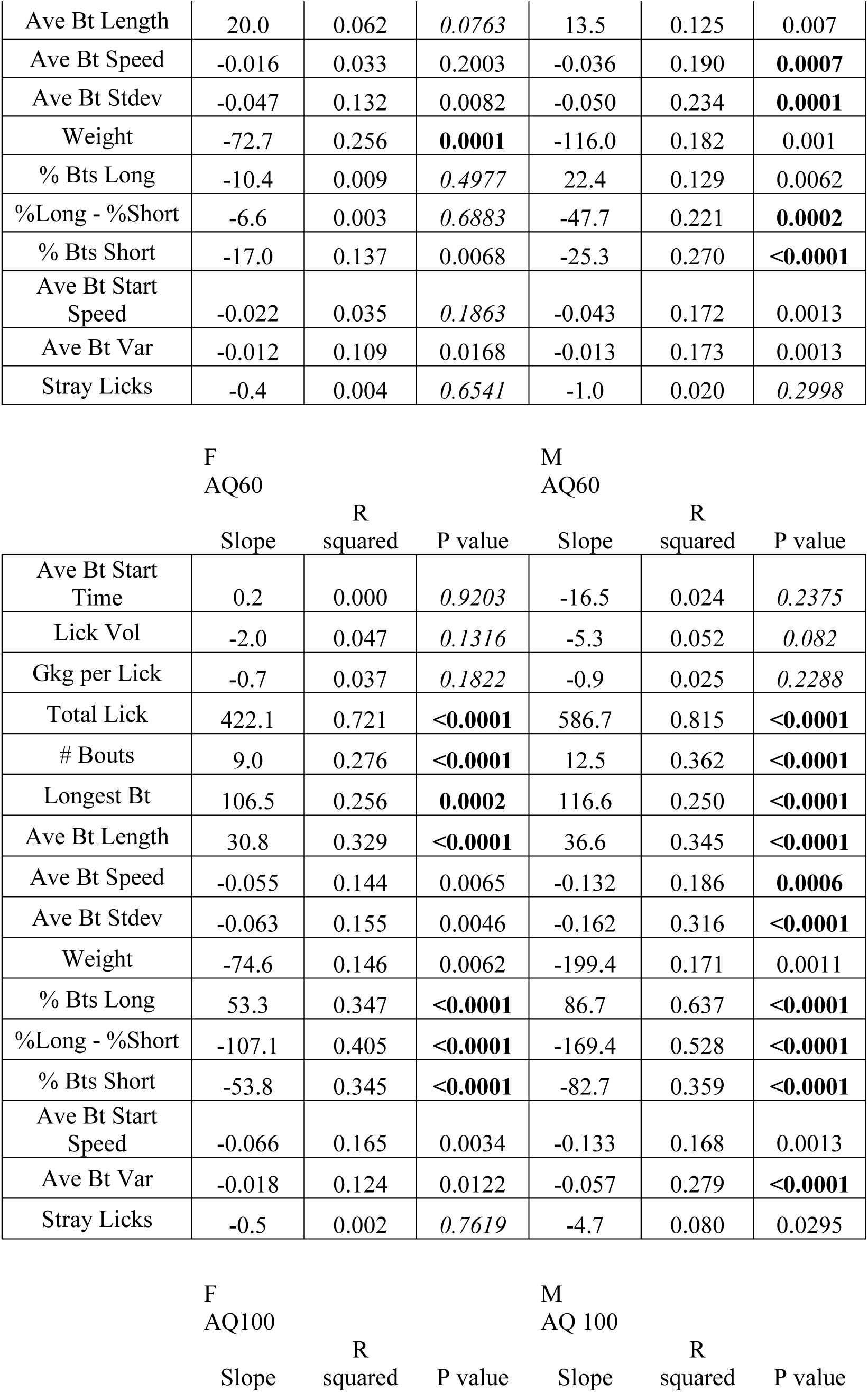
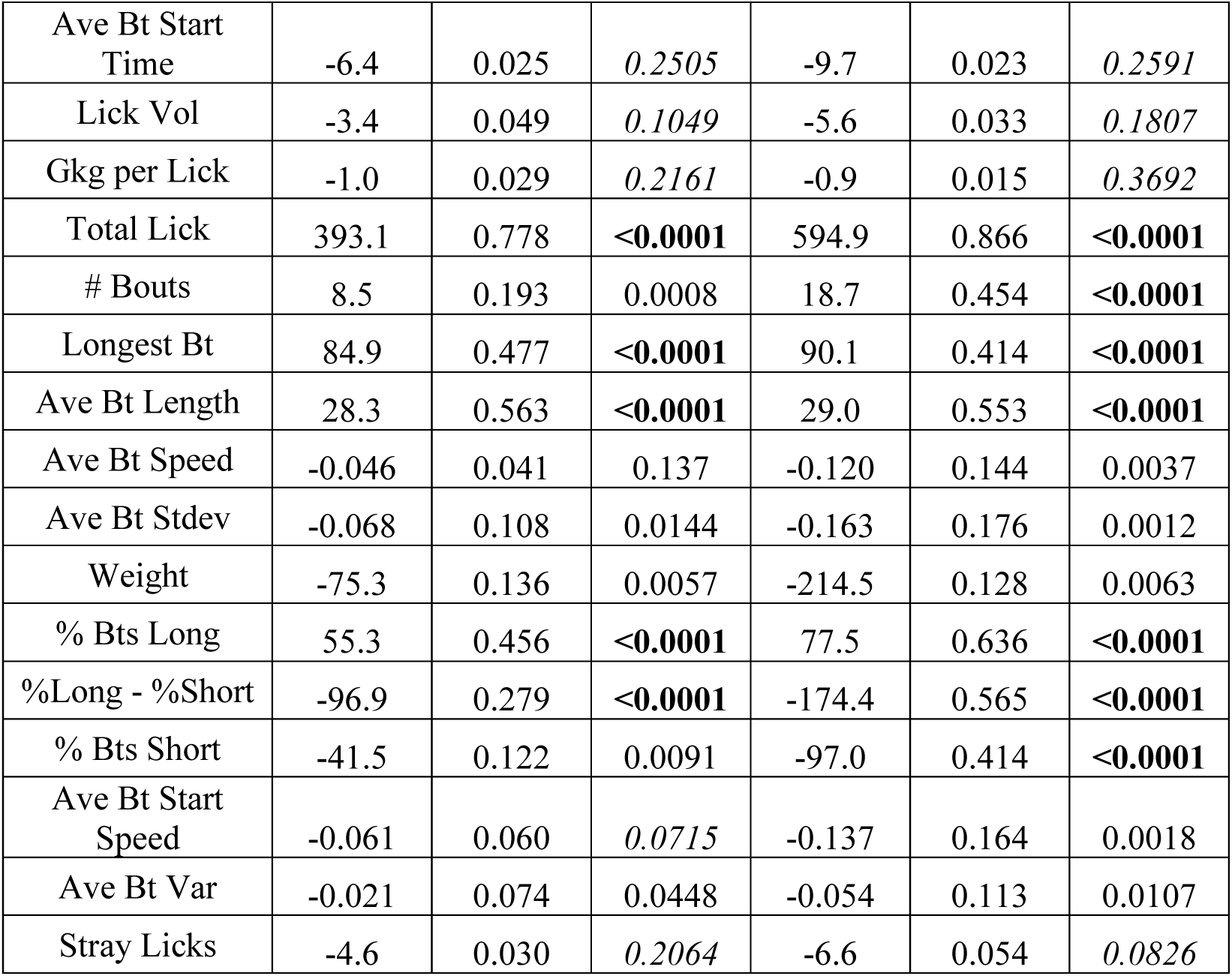
Alcohol intake (g/kg) correlated with other session measures. Since we consider total drinking level (in g/kg alcohol consumed) of primary importance, we performed a series of correlations to determine whether g/kg was significantly related to other session-level measures. With multiple corrections (considering the same 8 variables from the PCA, and 8 conditions), p<0.00078 is needed to be considered significant. This Table shows the p values for each sex and drinking condition. Bold shows significant, nonbold nonitalics is trend, and italics is non significant.

